# Physical activity in adolescence: cross-national comparisons of levels, distributions and disparities across 52 countries

**DOI:** 10.1101/483552

**Authors:** David Bann, Shaun Scholes, Meg Fluharty, Nikki Shure

## Abstract

**Introduction:** Despite global concerns regarding physical inactivity, limited cross-national evidence exists to compare adolescents’ physical activity participation. We analysed 52 high- and low-middle income countries, with activity undertaken inside and outside of school in 2015. We investigated gender- and socioeconomic-disparities, and additionally examined correlations with country-level indices of physical education (PE) curriculum time allocation, wealth, and income inequality.

**Methods:** We used the Programme for International Student Assessment (PISA), a nationally representative cross-sectional survey of 15-year-olds (N=347,935). Students reported average attendance (days/week) in PE classes, and the days/week engaged in moderate activity (MPA) and vigorous activity (VPA) outside of school. Both the mean and distributions of outcomes were evaluated, as were gender- and socioeconomic-disparities. Pearson’s correlations (r) between the physical activity outcomes and PE curriculum time allocation, wealth (indexed by GDP) and income inequality (indexed by the Gini coefficient) were calculated.

**Results:** Activity levels inside and outside of school were higher in Eastern Europe than Western Europe, the Americas, and the Middle East/North Africa. Comparisons of average levels masked potentially important differences in distributions. For example, activity levels inside school showed a bimodal distribution in the US (mean PE class attendance 2.4 days/week; 41.3%, 6.3% and 33.1% of students attended PE classes on 0, 2 and 5 days/week respectively). In contrast, most other countries exhibited more centrally shaped distributions. Pro-male and pro-high socioeconomic disparities were modest for participation inside school, but higher for MPA and VPA outside of school. The magnitude of these also differed markedly by country. Activity in school was weakly positively correlated with PE curriculum time allocation (r=0.33); activity outside of school was strongly negatively correlated with income inequality (e.g. r=-0.69 for MPA).

**Conclusion:** Our findings reveal extensive cross-country differences in adolescents’ physical activity; in turn, these highlight policy areas that could ultimately improve global adolescent health, such as the incorporation of minimum country-level PE classes, and the targeting of gender- and socioeconomic- disparities in activity conducted outside of school. Our findings also highlight the utility of educational databases such as PISA for use in global population health research.

## Introduction

Being physically active is widely thought to benefit mental, physical and social health,^1^ yet existing evidence suggests a global pandemic of physical inactivity. In 2010, more than 80% of school-going adolescents were estimated to be insufficiently physically active worldwide,^2 3^ yet substantial variation exists between countries.^4^ Documenting and understanding these between-country differences is important in order to identify countries and corresponding policies associated with particularly low or high levels of activity and enable benchmarking for future goal setting.

While there is some evidence that activity levels among adolescents are particularly high in northern European countries,^5 6^ previous cross-country comparisons of adolescent physical activity^5 7-12^ have so far produced limited evidence for a number of reasons. First, cross-national comparisons have been limited in geographic range, being primarily North American and/or European,^10 11 13^ and have under-represented low- and middle-income countries (LMICs) due to the lack of available surveillance data.^7 9^ Second, studies have not analysed activity performed inside and outside of school separately. Since both have different determinants, with modifiable educational policies potentially more important for physical activity undertaken in schools, it is likely to be useful to understand cross-country differences in each component separately. Third, studies have tended to compare countries using single numerical estimates of activity (e.g. averages or binary prevalence measures)—such comparisons may miss other important differences between countries in the distribution of activity outcomes. Fourth, not all studies have compared gender and socioeconomic status (SES) related disparities, which are additional policy concerns, further limiting the available evidence base.

Using a large-scale education achievement database to our knowledge previously unused in the epidemiological literature—The Programme for International Student Assessment (PISA)—we compared adolescent physical activity across 52 countries measured in 2015, spanning both high- and low-income countries, and activities undertaken both inside and outside of school. Since single numerical estimates may mask other meaningful cross-national differences, we also compared the distribution of each activity outcome, and additionally investigated gender- and SES-disparities. Finally, we additionally examined whether country-level physical education (PE) curriculum time allocation was correlated with the PISA assessed levels of activity participation inside school, and examined whether two structural factors thought to be key determinants of adolescent health^14^ —national levels of wealth and income inequality— were correlated with levels of activity both inside and outside of school.

## Methods

### Data source

PISA is conducted by the Organisation for Economic Co-operation and Development (OECD) in over 70 member and non-member nations and economies.^15^ PISA aims to draw a representative sample of in-school pupils in each country aged between 15 years and 3-months and 16 years and 2-months at the time of assessment. It has taken place every three years since 2000 yet physical activity data were included only in 2015. PISA has a two-stage probabilistic, stratified and clustered survey design. First, schools are stratified and then randomly selected with probability proportional to size (within a minimum of 150 schools from within each country). All countries and economies must ensure they meet the OECD’s response rate of 85% for schools and 80% for pupils in order to be included in the study^15^

Over half a million students participated in 2015, representing about 29 million students in the schools of the 72 participating countries and economies. To aid comparison, we restricted our analyses to 52 countries with available physical activity data: additional sub samples (‘economies’) were not included given concerns about national representation (e.g., the only four regions sampled in China were Beijing, Shanghai, Jiangsu, and Guangdong). We grouped countries into six regions: Eastern Europe; Western and Northern Europe; Asia; Middle East and North Africa; Americas; and Australasia. Further details of the PISA 2015 study are available elsewhere.^15^

### Physical activity

Students were asked to report *outside of school* the number of days during which they engaged in moderate physical activity (hereafter referred to as MPA: such as walking, climbing stairs or riding a bike to school) for at least 60 minutes per day (min/day) during the week before the PISA assessment. A similar question was asked for vigorous activity (hereafter referred to as VPA: such as running, cycling, aerobics, soccer and skating that makes you sweat and breathe hard) for at least 20 min/day. PISA also asked students, on average, on how many days they attended PE classes *during school* each week throughout the school year. Each outcome was summarised as days/week (range: 0-7).

### Socioeconomic status (SES)

SES was measured by reported family wealth possessions, a continuous variable estimated using item response theory scaling by the OECD. We calculated SES using eight standardised questions on possessions in and characteristics of the home. These included questions on whether the home has an internet connection, whether the student has her own room, the number of rooms in the home with a bath or shower, the number of televisions, computers, tablets, and e-book readers in the home, the number of cars the family has, and three country-specific wealth items (see^15^). Country-specific quintiles of this continuous variable were calculated for use in our analyses.

### Statistical Analysis

For each country, we calculated the mean number of days that students: (1) attended PE classes each week during the school year; (2) engaged in MPA in the last week (for ≥60min/day) outside of school; and (3) engaged in VPA in the last week (for ≥20min/day) outside of school. Using data aggregated at the country level, Pearson correlational analyses (r) were performed to verify the PISA data by comparing each indicator to the WHO 2010 compiled estimates of insufficient physical activity among school-going adolescents (aged 11-17 years) of both genders (% <60min/day of moderate- to vigorous-intensity activity).^16^ Students with missing data for gender, SES, and physical activity were excluded from analyses. To inform the potential for this in biasing our findings, logistic regression was used to examine demographic differences between students with and without physical activity data.

Cross-national comparisons were made by estimating mean (95% CI) activity levels within each country (days/week); these were calculated separately by gender and SES (top versus bottom wealth quintile given evidence for linearity) to examine disparities. We decided, a priori, to calculate wealth quintile specific estimates separately for male and female students due to expected gender differences as reported in the literature.^4 12^ Comparisons between countries’ physical activity distributions (e.g. the percentage of students active on 0, 1 and 2 days/week) were made by both tabulating and plotting via histograms.

Additional ecological analyses were conducted to examine factors plausibly correlated with—or be structural determinants of—cross-national differences in mean levels of activity. First, we examined Pearson correlation coefficients between PE class attendance and country-level PE curriculum time allocation for secondary schools (mean min/week) estimated by UNESCO in 2014.^17^ Second, we examined Pearson correlation coefficients between all activity outcomes and two economic outcomes collated by the World Bank—national wealth (as indexed by gross domestic product (GDP) per capita) and income inequality (as indexed by the Gini coefficient) in 2015 (or nearest neighbouring year to 2010 if not available in 2015).^18^

Analyses were performed using Stata V15.0 following the recommended use of the Balanced-Repeated-Replication (BRR) weights (final student response and replicate weights) to account for the amount of uncertainty due to sampling error, including the clustering of students within schools.^19^ Analytical syntax and accompanying datasets to enable replication of our findings are available at https://github.com/dbann/pisa.

## Results

Sample characteristics and descriptive statistics are summarised in Supplementary Table 1. Data on physical activity by gender was available for N=347,935 students, across 52 countries with an average (median) sample size of 5557 (range: 3150-18680). SES data was available for >99% of students (N=347,801); physical activity data was missing for 10% of students: lower family wealth, lower parental education, and being male were associated with increased odds of having missing data (P<0.001; data not shown). At the country level each PISA assessed activity outcome was inversely correlated with the WHO 2010 prevalence estimates of insufficient physical activity (PE classes: r=-0.14; MPA: r=-0.26; VPA: r=-0.29).

### Country differences in physical activity

There were large differences between regions in participation: activity levels inside and outside of school were highest in Eastern Europe; average activity levels outside of school were lowest in Middle East and North Africa (Figure 1; Supplementary Table 2). There were also notable differences within regions. For example, within Eastern Europe, average days/week in PE classes in Hungary were approximately double those in Croatia among both genders, whilst activity levels outside of school were higher by approximately 0.5 days/week.

**Figure 1:**
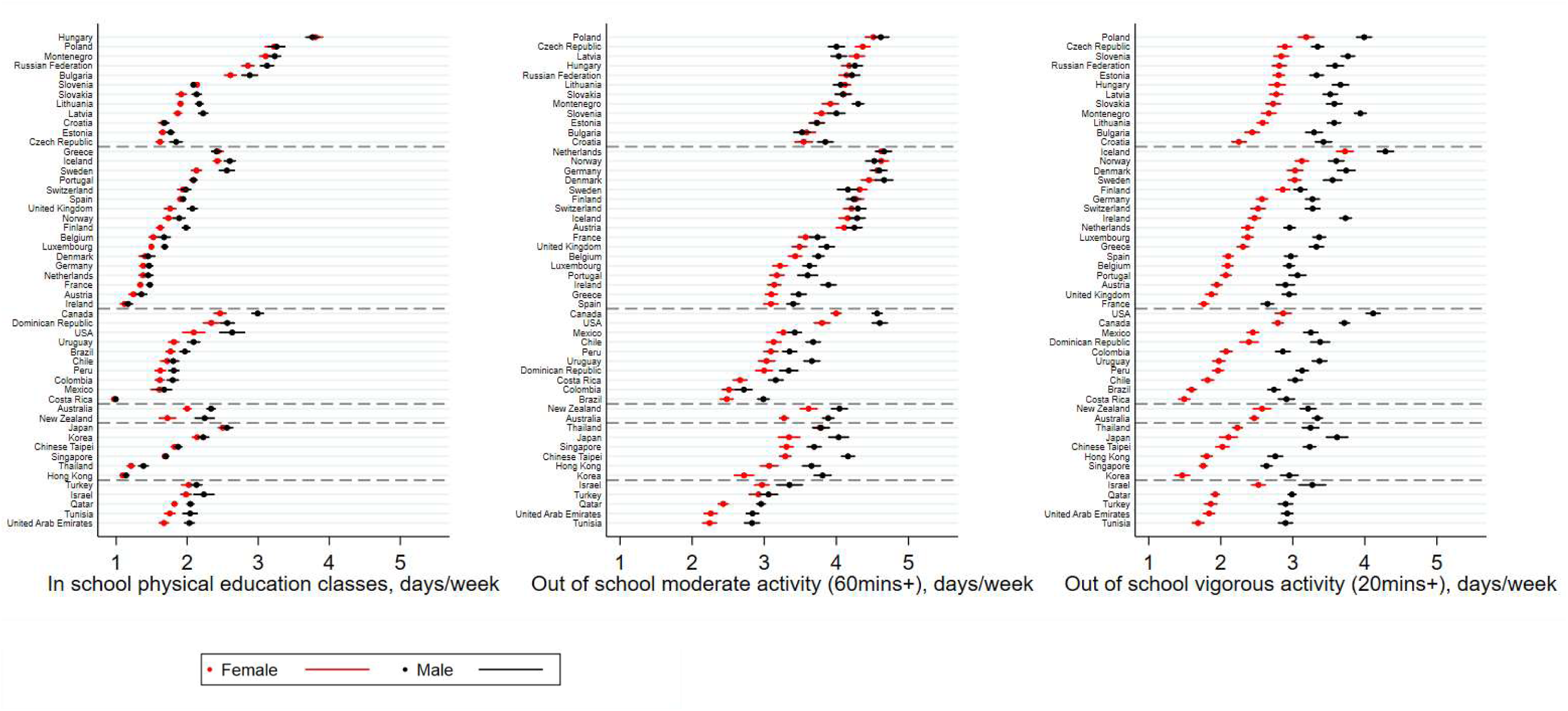
Gender disparities in adolescents’ mean (95% CI) physical activity: in school and out of school Note: reference period for school activity was in the last year; out of school was the last 7 days.

There was substantial diversity in the distribution of activity outcomes, particularly for activity inside school, revealing cross-country differences not found when using the mean as a single numerical summary measure (Figure 2; Supplementary Table 3). For example, activity levels inside school in the US showed a bimodal distribution (mean PE class attendance 2.3 days/week; 41.3%, 6.3% and 33.1% of students attended PE classes on 0, 2 and 5 days/week respectively), as did those in Canada. In contrast, most other countries exhibited more centrally shaped distributions (e.g. Sweden: mean 2.3 days/week; 2.0%, 64.3% and 1.8% of students attended PE classes on 0, 2 and 5 days/week respectively).

**Figure 2:**
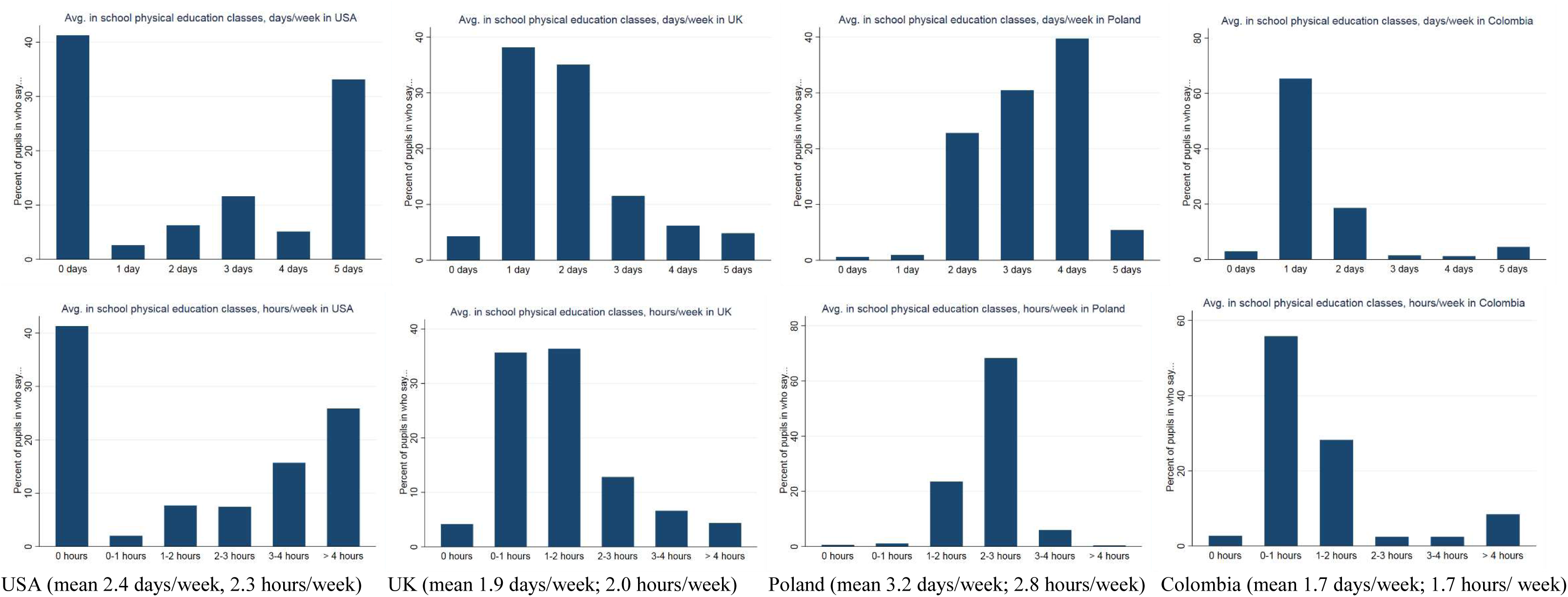
Histograms showing distributions of in-school physical activity participation in 4 countries with differing distributions.

### Gender and SES disparities

Males were more active than females, for activities inside and outside of school (Figure 1 and Supplementary Table 2 shows averages; Supplementary Table 4 shows distributions). Gender disparities however were most pronounced for activities outside of school and were larger for VPA (mean days/week: 3.4 males; 2.3 females) than for MPA (3.9 males; 3.4 females). Cross-national differences in gender disparities was most pronounced for VPA: being highest in the Americas and Asia (e.g. 1.4-1.5 days/week higher for males than females in Costa Rica, Uruguay, and Korea). Gender disparities for VPA were lower on average in Europe but there were outlying countries (e.g. 0.4 days/week higher for males than females in Finland; 1.3 days/week higher for males than females in Montenegro and in the Republic of Ireland). In some countries, gender disparities in average levels reflected differences at the upper tail of the distribution. For example, the average days/week engaged in MPA was 3.9 for males and 3.3 for females in Australia; 23.9% of males engaged in MPA seven days a week, while 13.2% of females did so.

SES disparities were largest for activities outside of school (Figures 3 and 4 show the averages for males and females respectively; Supplementary Table 5 shows the distributions). For both genders, activity levels for MPA and VPA were typically ~0.5 days/week higher for students in the top-versus bottom-wealth quintile. Regional variation was lower for SES- than for gender-disparities. However, the gap between the extreme wealth quintiles was higher than average among females for MPA in the Americas (e.g. 1 day/week higher for females in the top-versus bottom-quintile in the US) and among males for VPA in Eastern Europe (e.g. 1 day/week higher for males in the top-versus bottom-quintile in Lithuania). SES disparities in VPA were notably lower than the overall average in Asia among males and females.

**Figure 3:**
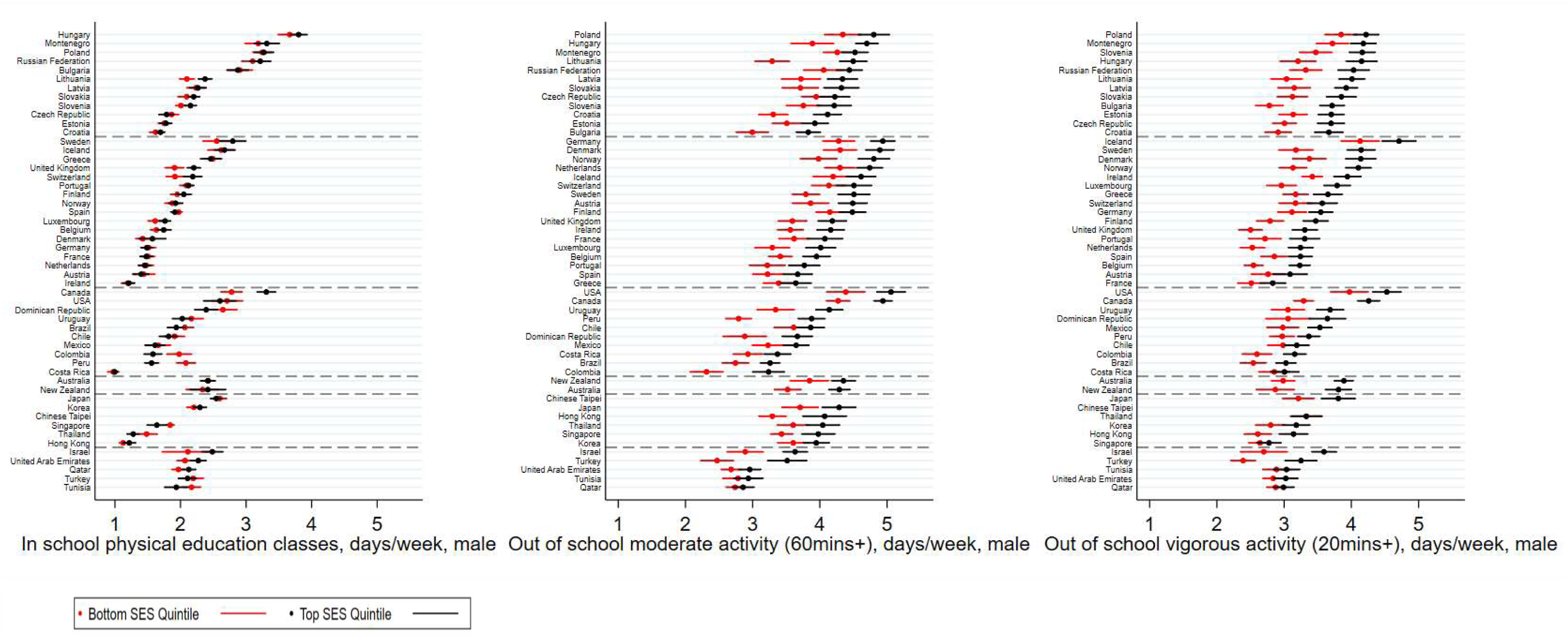
Socioeconomic (wealth-based) disparities in adolescents’ mean (95% CI) physical activity: in school and out of school among males Note: reference period for school activity was in the last year; out of school was the last 7 days.

**Figure 4:**
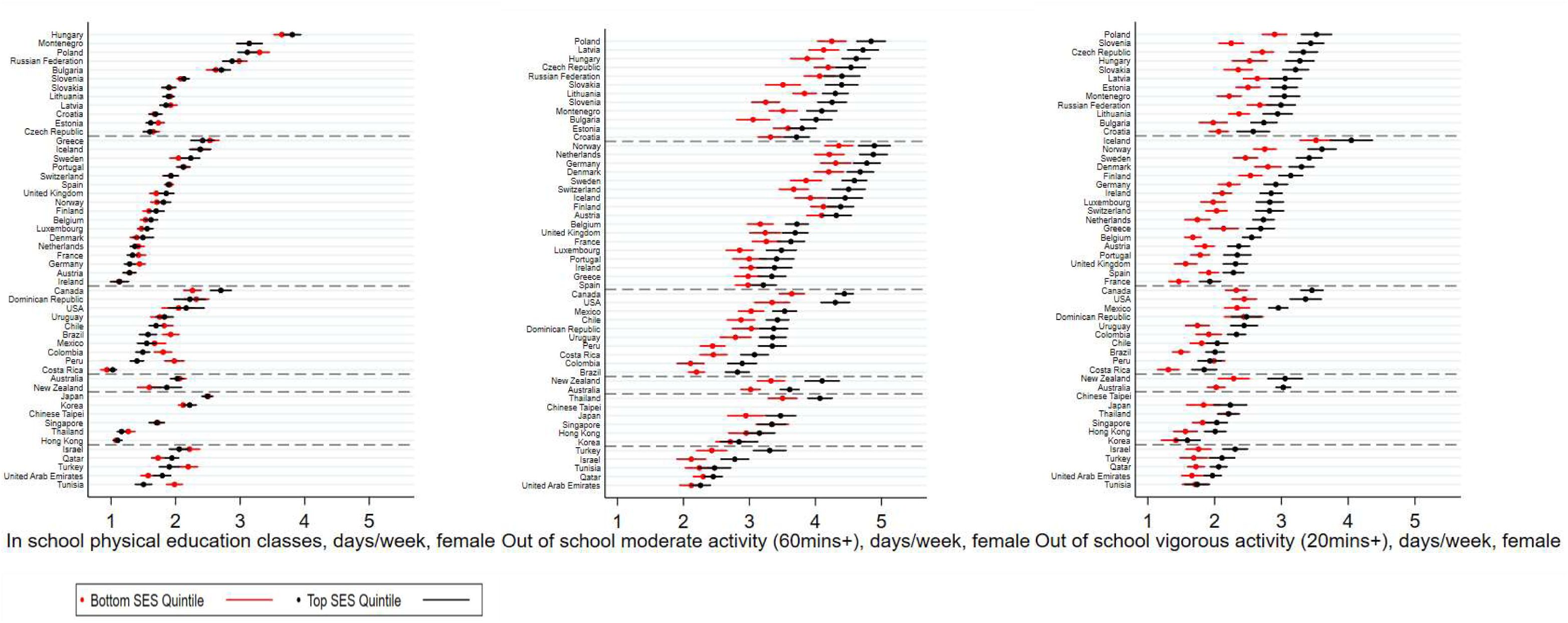
Socioeconomic (wealth-based) disparities in adolescents’ mean (95% CI) physical activity: in school and out of school among females. Note: reference period for school activity was in the last year; out of school was the last 7 days.

### Ecological analyses

Figures 5-6 show the results of the ecological analyses. Country-level PE curriculum time allocation for secondary schools (mean min/week) were positively correlated with levels of activity inside school (PE class attendance: r=0.23; number of days multiplied by average class time: r=0.33). National wealth as indexed by GDP was weakly negatively correlated with levels of activity inside school (r=-0.23), yet positively correlated with activity outside of school (MPA: r=0.41, VPA: r=0.26). Income inequality as indexed by the Gini coefficient was negatively correlated with each outcome, more strongly for levels of activity outside of school (MPA: r=-0.69, VPA r=-0.50) than for levels of activity inside school (r=-0.10).

**Figure 5:**
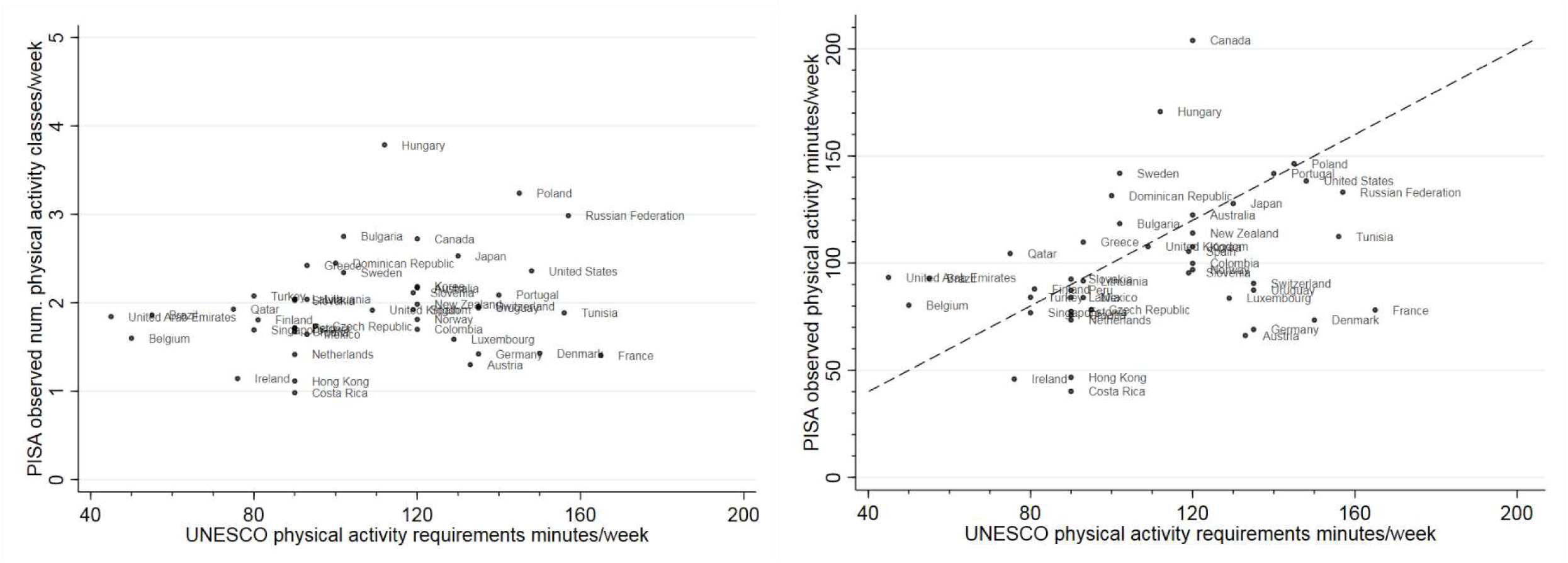
Scatter plots between country-level physical education (PE) curriculum time allocation and average in school physical activity levels

**Figure 6:**
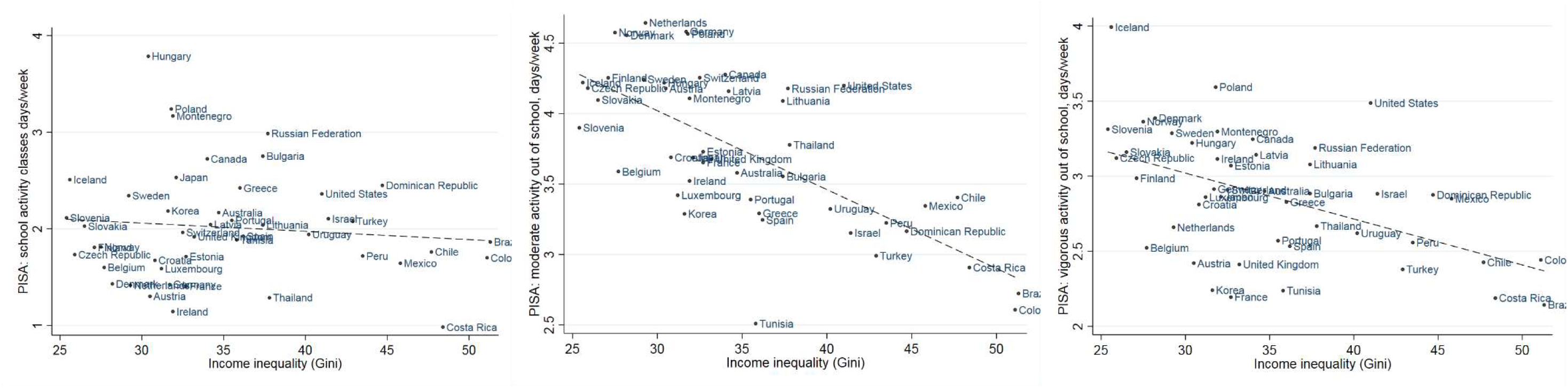
Scatter plots between country-level income inequality (Gini coefficient) and average physical activity levels

## Discussion

Using a large-scale global education database, we identified substantial cross-country differences in adolescents’ physical activity. Our findings extend those conducted in either high^10 11 13^ or low-middle^12^ income countries by including a greater number of countries across income levels, using more recent data (2015), and expanding on the comparisons across countries. We examined activities conducted inside and outside of school separately, compared the distributions of activity in addition to averages, quantified gender- and SES-disparities, and examined correlations with national-level economic factors thought to be key determinants of adolescent health.

There are several explanations for cross-country differences in adolescents’ physical activity, which if confirmed, should lead to multiple avenues for policy development. For activity conducted inside school, we anticipated that cross-country differences in laws or guidelines mandating PE class participation are likely to be a main source of variation. Our analyses partly support this suggestion, given the positive (albeit weak) correlation between the PISA assessed indicator (days/week attending PE classes throughout the school year) and the UNESCO compiled estimates of PE curriculum time allocation in secondary schools. For example, Hungary has reportedly higher levels of PE time allocation in secondary schools than Croatia (145 vs 90 min/week), consistent with our findings for these countries which showed the average days/week in PE classes to be twice as high. The large heterogeneity across US-states revealed by the UNESCO study (e.g., 30 min/week in Iowa versus 200 min/week in California),^17^ which we were not powered to investigate, is potentially partly reflected in our own finding in the US PISA sample of a bimodal distribution for activity in school (shown in Figure 2). Consistent with our findings, a recent US-wide study also found a bimodal distribution which persisted from 1991 to 2015: possibly reflecting the fewer opportunities for PE in high-poverty schools.^20^ Discrepancies between PE time allocation and observed levels of activity participation in schools in many countries, all those below the 45 degree line in Figure 5, may suggest that laws or guidelines are not being implemented sufficiently, thereby requiring action to redress. For example, Denmark’s 2016 Report Card on Physical Activity for Children and Youth suggested that high investment and government-led initiatives to support physical activity have not translated into higher observed physical activity levels.^21^ Other education policies which could explain country differences in activity include whether PE class length is enforced with mandatory minimum of minutes (e.g., 135 minutes/week in Poland,^22^ yet no mandatory minimums exist across England, Colombia, nor all of the USA); the funding, availability and quality of facilities within schools; and the training of PE teachers.^5^ The importance of these factors on cross-national differences in activity participation inside schools warrants future empirical investigation, yet is likely to be challenging given lack of consistent data across countries,^5^ and the possibility of reverse causality (since education policies may arise due to concerns about prior low physical activity levels which track across time). In addition to educational policies (and their implementation), other plausible explanations for differences between countries include social norms or cultural differences regarding the value of PE, particularly if time spent in PE classes is interpreted as being in competition with academic achievement.^23^

The cross-country differences in physical activity levels outside of school shown in our study are likely influenced by a greater range of determinants operating through several levels of influence (i.e. individual, social and built environmental, and policy). These include economic factors which partly determine the resources and the quality of the environments which facilitate participation, including the opportunities available for active transportation to and from school, and cultural factors, such as country-level beliefs regarding the importance of physical activity for health and personal/group identity. In support of the latter, activity levels outside of school were substantially higher in many Eastern European countries compared with their wealthier counterparts in Western Europe and Asia. Such cultural factors are difficult to measure, and are not readily explained by simple metrics of success in sports (e.g., per capita Olympic medal success is higher in Western Europe—see http://www.medalspercapita.com) but may be potentially fruitful targets for identification and modification to increase activity levels. Factors such as country-level economic development and income inequality are noted in highly cited papers as being crucial determinants of adolescent health^14^ yet to date have been inconsistently associated with cross-national differences in adolescent physical activity levels in previous studies.^5 10^ Our findings add to this evidence base. While we observed that national levels of income inequality strongly negatively correlated with levels of activity outside of school, it is unclear why this is the case. It remains speculative as to whether national levels of income inequality has a causal effect on activity participation (and if so, what factors mediate this effect) or if there are other factors such as those related to economic development-changing patterns of transportation, increased use of technology and urbanization^24^ which operate in such a way to result in a spurious association.^25^

While our study included more countries than previous studies, inclusion of other countries would expand the possibilities for cross-country comparison; these include low-middle income countries such as mainland China and India, which account for large fractions of the total adolescent population worldwide. We also included multiple activity outcomes, asked in identical form in each country; these enabled international comparisons of activity participation both inside and outside of school. Each outcome correlated in the expected direction with activity data aggregated by the WHO,^16^ despite measurement differences likely weakening such correlation (e.g., exact ages sampled, scale of measurement, and year of data collection). However, systematic differences in over- or under-reporting activity may exacerbate differences between countries, genders (e.g., if males tend to over-report more than females), and socioeconomic groups (e.g., if low SES groups over-report more than high SES groups). Evidence to date however on such patterns is inconclusive.^26^ Nevertheless, combinations of self-report and objective measures may be useful in future comparisons. While the sample framework for PISA is designed to enable national representation, as in all cross-country comparisons there may be between-country differences in unobserved factors which could confound our findings (e.g., differences in sample selection within each country, or time of year of measurement); as such, triangulation from other data sources may be useful. Further research within and between countries is also needed to examine the extent to which cross-country differences in adolescent physical activity (and their determinants) are different to those in other key life stages—childhood, adulthood, and older adult life.

In summary, our findings suggest substantial variation in adolescents’ activity across regions and countries. The presence of these differences suggests that the global pandemic of physical inactivity is not universal, and may be averted by understanding and adopting best-practices in different countries.

## Supporting information

## Acknowledgements

DB is supported by the Economic and Social Research Council (grant number ES/M001660/1) and The Academy of Medical Sciences / Wellcome Trust (“Springboard Health of the Public in 2040” award: HOP001/1025). The funders had no role in study design, data collection and analysis, decision to publish, or preparation of the manuscript.

