## Supplementary material for "Physical activity in adolescence: cross-national comparisons of levels, distributions and disparities across 52 countries"

Supplementary Table 1. Descriptive statistics for the countries included in the PISA sample.

| Country | N | GDP per capita in constant 2010 $USD | Gini coefficient, % | Urban residence, % | Obesity prevalence in adolescence, % (WHO, 2010) | Physical inactivity in adolescence, % (WHO, 2010) |
| --- | --- | --- | --- | --- | --- | --- |
| Australia | 12,908 | 54800.4 | 34.7 | 89.4 | 12.2 | 83.9 |
| Austria | 6,499 | 47958.4 | 30.5 | 66.0 | 8.4 | 75.4 |
| Belgium | 8,539 | 45068.3 | 27.7 | 97.9 | 6.9 | 82.5 |
| Brazil | 15,861 | 11322.1 | 51.3 | 85.7 | 10.4 | 86.7 |
| Bulgaria | 5,122 | 7612.0 | 37.4 | 73.9 | 10.3 | 74.1 |
| Canada | 18,680 | 50303.8 | 34.0 | 81.8 | 12.1 | 77.3 |
| Chile | 6,641 | 14907.1 | 47.7 | 89.5 | 14.8 | 85.2 |
| Chinese Taipei | 7,613 |  |  |  |  |  |
| Colombia | 10,988 | 7446.2 | 51.1 | 76.4 | 6.7 | 85.2 |
| Costa Rica | 5,616 | 9406.8 | 48.4 | 76.8 | 11.8 | 81.9 |
| Croatia | 5,439 | 13936.0 | 30.8 | 59.0 | 10.4 | 79.3 |
| Czech Republic | 6,463 | 21381.7 | 25.9 | 73.0 | 9.3 | 76.9 |
| Denmark | 6,350 | 59967.7 | 28.2 | 87.7 | 7.1 | 88.3 |
| Dominican Republic | 3,792 | 6552.7 | 44.7 | 79.0 | 14.5 |  |
| Estonia | 5,406 | 17734.0 | 32.7 | 67.5 | 6 | 86.0 |
| Finland | 5,580 | 45151.5 | 27.1 | 84.2 | 8.9 | 77.0 |
| France | 5,411 | 41689.7 | 32.7 | 79.5 | 7.9 | 88.1 |
| Germany | 4,846 | 45412.6 | 31.7 | 75.3 | 8.7 | 83.1 |
| Greece | 5,132 | 22648.8 | 36.0 | 78.0 | 13.5 | 85.9 |
| Hong Kong | 5,204 | 36262.9 |  | 100.0 |  |  |
| Hungary | 5,296 | 14629.2 | 30.4 | 71.2 | 10.6 | 80.8 |
| Iceland | 3,150 | 46221.9 | 25.6 | 94.1 | 9.8 | 83.8 |
| Ireland | 5,475 | 68030.8 | 31.9 | 63.2 | 9.5 |  |
| Israel | 6,265 | 32993.3 | 41.4 | 92.1 | 11.8 | 84.6 |
| Japan | 6,388 | 47163.5 | 32.1 | 93.5 | 3.3 |  |
| Korea | 5,499 | 24870.8 | 31.6 | 82.5 | 8.1 | 94.8 |
| Latvia | 4,667 | 14284.3 | 34.2 | 67.4 | 6.7 | 79.7 |
| Lithuania | 6,030 | 15383.5 | 37.4 | 66.5 | 6.5 | 83.4 |
| Luxembourg | 4,813 | 107648.6 | 31.2 | 90.2 | 8.2 |  |
| Mexico | 7,006 | 9615.3 | 45.8 | 79.2 | 14.4 |  |
| Montenegro | 4,789 | 7279.9 | 31.9 | 64.0 | 7.2 |  |
| Netherlands | 5,012 | 51410.5 | 29.3 | 90.5 | 6.9 | 81.1 |
| New Zealand | 4,100 | 36191.5 |  | 86.3 | 16 |  |
| Norway | 4,982 | 90104.0 | 27.5 | 80.5 | 8.9 | 85.0 |
| Peru | 6,343 | 5936.0 | 43.5 | 78.6 | 7.5 | 84.9 |
| Poland | 4,318 | 14640.2 | 31.8 | 60.5 | 8.6 | 79.6 |
| Portugal | 6,794 | 22016.8 | 35.5 | 63.5 | 10.3 | 86.7 |
| Qatar | 10,510 | 67277.2 |  | 99.2 | 19.2 | 90.1 |
| Russian Federation | 5,513 | 11325.8 | 37.7 | 74.0 | 6.7 | 87.5 |
| Singapore | 5,968 | 52244.6 |  | 100.0 | 6.7 | 89.9 |
| Slovakia | 5,875 | 18678.9 | 26.5 | 53.6 | 7.6 | 77.4 |
| Slovenia | 6,008 | 23731.2 | 25.4 | 49.7 | 8.7 | 79.8 |
| Spain | 6,477 | 30532.4 | 36.2 | 79.6 | 10.6 | 77.2 |
| Sweden | 4,943 | 55395.1 | 29.2 | 85.8 | 6.6 | 86.1 |
| Switzerland | 5,403 | 76472.5 | 32.5 | 73.9 | 5.6 | 88.0 |
| Thailand | 7,658 | 5733.9 | 37.8 | 50.4 | 10.7 | 84.4 |
| Tunisia | 4,650 | 4264.5 | 35.8 | 66.8 | 8.1 | 81.4 |
| Turkey | 5,534 | 13898.3 | 42.9 | 73.4 | 11.1 | 82.2 |
| United Arab Emirates | 13,061 | 40159.6 |  | 85.5 | 16.7 | 82.6 |
| United Kingdom | 12,909 | 41536.9 | 33.2 | 82.6 | 10.1 | 79.0 |
| United States | 5,294 | 51956.6 | 41.0 | 81.6 | 21.2 | 72.6 |
| Uruguay | 5,115 | 13859.4 | 40.2 | 95.3 | 13.5 |  |

Note: missing cells shown where data were not published by the source: World Bank (Gini, GDP, urban residence) or WHO (obesity and physical inactivity)

Supplementary Table 2. Mean physical activity outcomes by country (overall, by gender, and by wealth for each gender)

| **(a) Physical activity in school, classes per week** | | | | | | | | | | | |  |
| --- | --- | --- | --- | --- | --- | --- | --- | --- | --- | --- | --- | --- |
|  | Overall |  |  |  | Female |  |  |  | Male |  |  |  |
| Country | Mean | Median | SD | N | Mean | Median | SD | N | Mean | Median | SD | N |
| Australia | 2.2 | 2 | 1.4 | 12908 | 2.0 | 2 | 1.4 | 6459 | 2.3 | 2 | 1.5 | 6449 |
| Austria | 1.3 | 1 | 1.1 | 6499 | 1.2 | 1 | 0.9 | 3236 | 1.4 | 1 | 1.2 | 3263 |
| Belgium | 1.6 | 1 | 0.9 | 8539 | 1.5 | 1 | 0.8 | 4246 | 1.7 | 1 | 1.0 | 4293 |
| Brazil | 1.9 | 1 | 1.7 | 15861 | 1.8 | 1 | 1.6 | 8372 | 2.0 | 2 | 1.7 | 7489 |
| Bulgaria | 2.8 | 2 | 1.5 | 5122 | 2.6 | 2 | 1.3 | 2466 | 2.9 | 2 | 1.6 | 2656 |
| Canada | 2.7 | 3 | 2.0 | 18680 | 2.5 | 2 | 2.0 | 9440 | 3.0 | 3 | 2.0 | 9240 |
| Chile | 1.8 | 1 | 1.2 | 6641 | 1.7 | 1 | 1.2 | 3305 | 1.8 | 1 | 1.2 | 3336 |
| Chinese Taipei | 1.8 | 2 | 0.7 | 7613 | 1.8 | 2 | 0.6 | 3775 | 1.9 | 2 | 0.7 | 3838 |
| Colombia | 1.7 | 1 | 1.5 | 10988 | 1.6 | 1 | 1.4 | 5771 | 1.8 | 1 | 1.5 | 5217 |
| Costa Rica | 1.0 | 1 | 0.5 | 5616 | 1.0 | 1 | 0.4 | 2876 | 1.0 | 1 | 0.6 | 2740 |
| Croatia | 1.7 | 2 | 0.7 | 5439 | 1.7 | 2 | 0.6 | 2848 | 1.7 | 2 | 0.7 | 2591 |
| Czech Republic | 1.7 | 2 | 0.9 | 6463 | 1.6 | 2 | 0.8 | 3242 | 1.8 | 2 | 1.0 | 3221 |
| Denmark | 1.4 | 1 | 1.0 | 6350 | 1.4 | 1 | 1.0 | 3230 | 1.4 | 1 | 1.0 | 3120 |
| Dominican Republic | 2.4 | 2 | 1.7 | 3792 | 2.3 | 2 | 1.6 | 1989 | 2.6 | 2 | 1.7 | 1803 |
| Estonia | 1.7 | 2 | 0.8 | 5406 | 1.7 | 2 | 0.7 | 2727 | 1.8 | 2 | 0.9 | 2679 |
| Finland | 1.8 | 2 | 1.1 | 5580 | 1.6 | 1 | 1.0 | 2753 | 2.0 | 2 | 1.2 | 2827 |
| France | 1.4 | 1 | 1.0 | 5411 | 1.3 | 1 | 0.9 | 2811 | 1.5 | 1 | 1.0 | 2600 |
| Germany | 1.4 | 1 | 0.9 | 4846 | 1.4 | 1 | 0.8 | 2482 | 1.5 | 1 | 0.9 | 2364 |
| Greece | 2.4 | 2 | 1.5 | 5132 | 2.4 | 2 | 1.6 | 2558 | 2.4 | 2 | 1.5 | 2574 |
| Hong Kong | 1.1 | 1 | 0.5 | 5204 | 1.1 | 1 | 0.5 | 2610 | 1.1 | 1 | 0.6 | 2594 |
| Hungary | 3.8 | 4 | 1.2 | 5296 | 3.8 | 4 | 1.2 | 2684 | 3.8 | 4 | 1.2 | 2612 |
| Iceland | 2.5 | 2 | 1.5 | 3150 | 2.4 | 2 | 1.4 | 1648 | 2.6 | 2 | 1.5 | 1502 |
| Ireland | 1.1 | 1 | 0.8 | 5475 | 1.1 | 1 | 0.7 | 2713 | 1.2 | 1 | 0.9 | 2762 |
| Israel | 2.1 | 2 | 1.7 | 6265 | 2.0 | 2 | 1.5 | 3535 | 2.2 | 2 | 1.9 | 2730 |
| Japan | 2.5 | 3 | 0.7 | 6388 | 2.5 | 3 | 0.6 | 3199 | 2.6 | 3 | 0.7 | 3189 |
| Korea | 2.2 | 2 | 0.7 | 5499 | 2.1 | 2 | 0.6 | 2633 | 2.2 | 2 | 0.8 | 2866 |
| Latvia | 2.0 | 2 | 1.0 | 4667 | 1.9 | 2 | 0.9 | 2345 | 2.2 | 2 | 1.1 | 2322 |
| Lithuania | 2.0 | 2 | 1.0 | 6030 | 1.9 | 2 | 0.8 | 3005 | 2.2 | 2 | 1.1 | 3025 |
| Luxembourg | 1.6 | 1 | 1.0 | 4813 | 1.5 | 1 | 0.9 | 2462 | 1.7 | 1 | 1.1 | 2351 |
| Mexico | 1.6 | 1 | 1.5 | 7006 | 1.6 | 1 | 1.5 | 3550 | 1.7 | 2 | 1.5 | 3456 |
| Montenegro | 3.2 | 2 | 2.1 | 4789 | 3.1 | 2 | 2.1 | 2415 | 3.2 | 2 | 2.1 | 2374 |
| Netherlands | 1.4 | 1 | 0.8 | 5012 | 1.4 | 1 | 0.7 | 2545 | 1.5 | 1 | 0.8 | 2467 |
| New Zealand | 2.0 | 2 | 1.9 | 4100 | 1.7 | 1 | 1.9 | 2062 | 2.2 | 2 | 2.0 | 2038 |
| Norway | 1.8 | 2 | 0.9 | 4982 | 1.7 | 2 | 0.8 | 2512 | 1.9 | 2 | 1.0 | 2470 |
| Peru | 1.7 | 1 | 1.4 | 6343 | 1.6 | 1 | 1.3 | 3099 | 1.8 | 1 | 1.5 | 3244 |
| Poland | 3.2 | 3 | 0.9 | 4318 | 3.2 | 3 | 0.9 | 2148 | 3.3 | 3 | 0.9 | 2170 |
| Portugal | 2.1 | 2 | 0.9 | 6794 | 2.1 | 2 | 0.8 | 3424 | 2.1 | 2 | 0.9 | 3370 |
| Qatar | 1.9 | 1 | 1.6 | 10510 | 1.8 | 1 | 1.5 | 5671 | 2.0 | 1 | 1.6 | 4839 |
| Russian Federation | 3.0 | 3 | 1.3 | 5513 | 2.9 | 3 | 1.2 | 2872 | 3.1 | 3 | 1.4 | 2641 |
| Singapore | 1.7 | 2 | 0.8 | 5968 | 1.7 | 2 | 0.8 | 2909 | 1.7 | 2 | 0.8 | 3059 |
| Slovakia | 2.0 | 2 | 0.9 | 5875 | 1.9 | 2 | 0.8 | 2833 | 2.1 | 2 | 1.0 | 3042 |
| Slovenia | 2.1 | 2 | 0.9 | 6008 | 2.1 | 2 | 0.8 | 2779 | 2.1 | 2 | 0.9 | 3229 |
| Spain | 1.9 | 2 | 0.5 | 6477 | 1.9 | 2 | 0.5 | 3297 | 1.9 | 2 | 0.6 | 3180 |
| Sweden | 2.3 | 2 | 1.5 | 4943 | 2.1 | 2 | 1.3 | 2520 | 2.6 | 2 | 1.7 | 2423 |
| Switzerland | 2.0 | 2 | 1.0 | 5403 | 1.9 | 2 | 0.9 | 2609 | 2.0 | 2 | 1.1 | 2794 |
| Thailand | 1.3 | 1 | 1.0 | 7658 | 1.2 | 1 | 0.9 | 4319 | 1.4 | 1 | 1.2 | 3339 |
| Tunisia | 1.9 | 2 | 1.3 | 4650 | 1.8 | 2 | 1.2 | 2597 | 2.0 | 2 | 1.4 | 2053 |
| Turkey | 2.1 | 1 | 1.6 | 5534 | 2.0 | 1 | 1.6 | 2773 | 2.1 | 1 | 1.6 | 2761 |
| United Arab Emirates | 1.8 | 2 | 1.4 | 13061 | 1.7 | 1 | 1.2 | 6786 | 2.0 | 2 | 1.5 | 6275 |
| United Kingdom | 1.9 | 2 | 1.2 | 12909 | 1.8 | 2 | 1.1 | 6396 | 2.1 | 2 | 1.2 | 6513 |
| United States | 2.4 | 2 | 2.2 | 5294 | 2.1 | 2 | 2.2 | 2668 | 2.6 | 3 | 2.2 | 2626 |
| Uruguay | 1.9 | 2 | 1.5 | 5115 | 1.8 | 2 | 1.4 | 2719 | 2.1 | 2 | 1.6 | 2396 |

| **(b) Out of school moderate activity (60mins+), days/week** | | | | | | | | | | | |  |
| --- | --- | --- | --- | --- | --- | --- | --- | --- | --- | --- | --- | --- |
|  | Overall |  |  |  | Female |  |  |  | Male |  |  |  |
| Country | Mean | Median | SD | N | Mean | Median | SD | N | Mean | Median | SD | N |
| Australia | 3.6 | 3 | 2.3 | 12908 | 3.3 | 3 | 2.2 | 6459 | 3.9 | 4 | 2.4 | 6449 |
| Austria | 4.2 | 5 | 2.6 | 6499 | 4.1 | 4 | 2.5 | 3236 | 4.2 | 5 | 2.6 | 3263 |
| Belgium | 3.6 | 3 | 2.5 | 8539 | 3.4 | 3 | 2.5 | 4246 | 3.7 | 4 | 2.5 | 4293 |
| Brazil | 2.7 | 2 | 2.4 | 15861 | 2.5 | 2 | 2.4 | 8372 | 3.0 | 2 | 2.4 | 7489 |
| Bulgaria | 3.6 | 3 | 2.4 | 5122 | 3.6 | 3 | 2.4 | 2466 | 3.5 | 3 | 2.4 | 2656 |
| Canada | 4.3 | 5 | 2.3 | 18680 | 4.0 | 4 | 2.3 | 9440 | 4.6 | 5 | 2.3 | 9240 |
| Chile | 3.4 | 3 | 2.4 | 6641 | 3.1 | 3 | 2.4 | 3305 | 3.7 | 3 | 2.4 | 3336 |
| Chinese Taipei | 3.7 | 4 | 2.6 | 7613 | 3.3 | 3 | 2.6 | 3775 | 4.2 | 5 | 2.6 | 3838 |
| Colombia | 2.6 | 2 | 2.4 | 10988 | 2.5 | 2 | 2.4 | 5771 | 2.7 | 2 | 2.4 | 5217 |
| Costa Rica | 2.9 | 2 | 2.3 | 5616 | 2.7 | 2 | 2.2 | 2876 | 3.2 | 3 | 2.3 | 2740 |
| Croatia | 3.7 | 3 | 2.5 | 5439 | 3.5 | 3 | 2.5 | 2848 | 3.8 | 4 | 2.5 | 2591 |
| Czech Republic | 4.2 | 4 | 2.4 | 6463 | 4.4 | 5 | 2.4 | 3242 | 4.0 | 4 | 2.5 | 3221 |
| Denmark | 4.6 | 5 | 2.3 | 6350 | 4.5 | 5 | 2.3 | 3230 | 4.7 | 5 | 2.3 | 3120 |
| Dominican Republic | 3.2 | 3 | 2.4 | 3792 | 3.0 | 2 | 2.4 | 1989 | 3.3 | 3 | 2.4 | 1803 |
| Estonia | 3.7 | 4 | 2.3 | 5406 | 3.7 | 4 | 2.3 | 2727 | 3.7 | 4 | 2.3 | 2679 |
| Finland | 4.3 | 4 | 2.2 | 5580 | 4.3 | 4 | 2.2 | 2753 | 4.2 | 5 | 2.3 | 2827 |
| France | 3.7 | 3 | 2.6 | 5411 | 3.6 | 3 | 2.6 | 2811 | 3.7 | 4 | 2.5 | 2600 |
| Germany | 4.6 | 5 | 2.4 | 4846 | 4.6 | 5 | 2.3 | 2482 | 4.6 | 5 | 2.4 | 2364 |
| Greece | 3.3 | 3 | 2.4 | 5132 | 3.1 | 3 | 2.3 | 2558 | 3.5 | 3 | 2.4 | 2574 |
| Hong Kong | 3.4 | 3 | 2.6 | 5204 | 3.1 | 2 | 2.5 | 2610 | 3.7 | 3 | 2.6 | 2594 |
| Hungary | 4.2 | 5 | 2.4 | 5296 | 4.2 | 4 | 2.4 | 2684 | 4.3 | 5 | 2.4 | 2612 |
| Iceland | 4.2 | 5 | 2.4 | 3150 | 4.2 | 4 | 2.3 | 1648 | 4.3 | 5 | 2.4 | 1502 |
| Ireland | 3.5 | 3 | 2.3 | 5475 | 3.1 | 3 | 2.2 | 2713 | 3.9 | 4 | 2.3 | 2762 |
| Israel | 3.2 | 3 | 2.5 | 6265 | 3.0 | 2 | 2.5 | 3535 | 3.4 | 3 | 2.4 | 2730 |
| Japan | 3.7 | 5 | 2.8 | 6388 | 3.3 | 4 | 2.8 | 3199 | 4.0 | 5 | 2.8 | 3189 |
| Korea | 3.3 | 3 | 2.6 | 5499 | 2.7 | 2 | 2.5 | 2633 | 3.8 | 4 | 2.5 | 2866 |
| Latvia | 4.2 | 4 | 2.4 | 4667 | 4.3 | 4 | 2.3 | 2345 | 4.0 | 4 | 2.4 | 2322 |
| Lithuania | 4.1 | 4 | 2.4 | 6030 | 4.1 | 4 | 2.4 | 3005 | 4.1 | 4 | 2.5 | 3025 |
| Luxembourg | 3.4 | 3 | 2.4 | 4813 | 3.2 | 3 | 2.4 | 2462 | 3.6 | 3 | 2.4 | 2351 |
| Mexico | 3.3 | 3 | 2.3 | 7006 | 3.3 | 3 | 2.3 | 3550 | 3.4 | 3 | 2.3 | 3456 |
| Montenegro | 4.1 | 4 | 2.3 | 4789 | 3.9 | 4 | 2.3 | 2415 | 4.3 | 4 | 2.3 | 2374 |
| Netherlands | 4.6 | 5 | 2.2 | 5012 | 4.6 | 5 | 2.2 | 2545 | 4.7 | 5 | 2.2 | 2467 |
| New Zealand | 3.8 | 4 | 2.4 | 4100 | 3.6 | 3 | 2.3 | 2062 | 4.0 | 4 | 2.4 | 2038 |
| Norway | 4.6 | 5 | 2.3 | 4982 | 4.6 | 5 | 2.3 | 2512 | 4.5 | 5 | 2.4 | 2470 |
| Peru | 3.2 | 3 | 2.3 | 6343 | 3.1 | 3 | 2.3 | 3099 | 3.4 | 3 | 2.3 | 3244 |
| Poland | 4.6 | 5 | 2.4 | 4318 | 4.5 | 5 | 2.4 | 2148 | 4.6 | 5 | 2.4 | 2170 |
| Portugal | 3.4 | 3 | 2.5 | 6794 | 3.2 | 3 | 2.5 | 3424 | 3.6 | 3 | 2.6 | 3370 |
| Qatar | 2.7 | 2 | 2.4 | 10510 | 2.4 | 2 | 2.3 | 5671 | 3.0 | 3 | 2.4 | 4839 |
| Russian Federation | 4.2 | 4 | 2.3 | 5513 | 4.1 | 4 | 2.3 | 2872 | 4.2 | 4 | 2.3 | 2641 |
| Singapore | 3.5 | 3 | 2.6 | 5968 | 3.3 | 3 | 2.6 | 2909 | 3.7 | 3 | 2.6 | 3059 |
| Slovakia | 4.1 | 4 | 2.4 | 5875 | 4.1 | 4 | 2.4 | 2833 | 4.1 | 4 | 2.5 | 3042 |
| Slovenia | 3.9 | 4 | 2.3 | 6008 | 3.8 | 3 | 2.3 | 2779 | 4.0 | 4 | 2.4 | 3229 |
| Spain | 3.2 | 3 | 2.4 | 6477 | 3.1 | 3 | 2.3 | 3297 | 3.4 | 3 | 2.5 | 3180 |
| Sweden | 4.2 | 5 | 2.4 | 4943 | 4.3 | 5 | 2.4 | 2520 | 4.2 | 5 | 2.5 | 2423 |
| Switzerland | 4.3 | 5 | 2.5 | 5403 | 4.2 | 5 | 2.5 | 2609 | 4.3 | 5 | 2.4 | 2794 |
| Thailand | 3.8 | 3 | 2.3 | 7658 | 3.8 | 3 | 2.3 | 4319 | 3.8 | 3 | 2.4 | 3339 |
| Tunisia | 2.5 | 2 | 2.2 | 4650 | 2.2 | 2 | 2.2 | 2597 | 2.8 | 2 | 2.2 | 2053 |
| Turkey | 3.0 | 2 | 2.5 | 5534 | 2.9 | 2 | 2.5 | 2773 | 3.1 | 2 | 2.5 | 2761 |
| United Arab Emirates | 2.5 | 2 | 2.4 | 13061 | 2.3 | 2 | 2.3 | 6786 | 2.8 | 2 | 2.4 | 6275 |
| United Kingdom | 3.7 | 3 | 2.5 | 12909 | 3.5 | 3 | 2.5 | 6396 | 3.9 | 4 | 2.5 | 6513 |
| United States | 4.2 | 5 | 2.4 | 5294 | 3.8 | 4 | 2.4 | 2668 | 4.6 | 5 | 2.4 | 2626 |
| Uruguay | 3.3 | 3 | 2.4 | 5115 | 3.0 | 3 | 2.4 | 2719 | 3.7 | 3 | 2.4 | 2396 |

| **(c) Out of school vigorous activity (20mins+), days/week** | | | |  |  |  |  |  |  |  |  |  |
| --- | --- | --- | --- | --- | --- | --- | --- | --- | --- | --- | --- | --- |
|  | Overall |  |  |  | Female |  |  |  | Male |  |  |  |
| Country | Mean | Median | SD | N | Mean | Median | SD | N | Mean | Median | SD | N |
| Australia | 2.9 | 3 | 2.2 | 12908 | 2.5 | 2 | 2.0 | 6459 | 3.3 | 3 | 2.3 | 6449 |
| Austria | 2.4 | 2 | 2.1 | 6499 | 1.9 | 2 | 1.8 | 3236 | 2.9 | 3 | 2.2 | 3263 |
| Belgium | 2.5 | 2 | 2.1 | 8539 | 2.1 | 2 | 1.9 | 4246 | 2.9 | 3 | 2.1 | 4293 |
| Brazil | 2.1 | 1 | 2.3 | 15861 | 1.6 | 1 | 2.1 | 8372 | 2.7 | 2 | 2.4 | 7489 |
| Bulgaria | 2.9 | 2 | 2.3 | 5122 | 2.4 | 2 | 2.1 | 2466 | 3.3 | 3 | 2.3 | 2656 |
| Canada | 3.2 | 3 | 2.3 | 18680 | 2.8 | 3 | 2.1 | 9440 | 3.7 | 4 | 2.3 | 9240 |
| Chile | 2.4 | 2 | 2.1 | 6641 | 1.8 | 1 | 1.9 | 3305 | 3.0 | 3 | 2.2 | 3336 |
| Chinese Taipei | 2.6 | 2 | 2.2 | 7613 | 2.0 | 2 | 1.9 | 3775 | 3.2 | 3 | 2.3 | 3838 |
| Colombia | 2.4 | 2 | 2.1 | 10988 | 2.1 | 1 | 2.0 | 5771 | 2.9 | 2 | 2.2 | 5217 |
| Costa Rica | 2.2 | 2 | 2.1 | 5616 | 1.5 | 1 | 1.8 | 2876 | 2.9 | 3 | 2.2 | 2740 |
| Croatia | 2.8 | 2 | 2.3 | 5439 | 2.3 | 2 | 2.1 | 2848 | 3.4 | 3 | 2.3 | 2591 |
| Czech Republic | 3.1 | 3 | 2.2 | 6463 | 2.9 | 3 | 2.1 | 3242 | 3.3 | 3 | 2.2 | 3221 |
| Denmark | 3.4 | 3 | 2.2 | 6350 | 3.0 | 3 | 2.1 | 3230 | 3.7 | 4 | 2.2 | 3120 |
| Dominican Republic | 2.9 | 2 | 2.3 | 3792 | 2.4 | 2 | 2.1 | 1989 | 3.4 | 3 | 2.3 | 1803 |
| Estonia | 3.1 | 3 | 2.1 | 5406 | 2.8 | 3 | 2.1 | 2727 | 3.3 | 3 | 2.2 | 2679 |
| Finland | 3.0 | 3 | 2.1 | 5580 | 2.9 | 3 | 1.9 | 2753 | 3.1 | 3 | 2.1 | 2827 |
| France | 2.2 | 2 | 2.0 | 5411 | 1.8 | 1 | 1.8 | 2811 | 2.6 | 2 | 2.1 | 2600 |
| Germany | 2.9 | 3 | 2.0 | 4846 | 2.6 | 2 | 1.8 | 2482 | 3.3 | 3 | 2.1 | 2364 |
| Greece | 2.8 | 3 | 2.2 | 5132 | 2.3 | 2 | 2.1 | 2558 | 3.3 | 3 | 2.3 | 2574 |
| Hong Kong | 2.3 | 2 | 2.1 | 5204 | 1.8 | 1 | 1.8 | 2610 | 2.8 | 2 | 2.3 | 2594 |
| Hungary | 3.2 | 3 | 2.2 | 5296 | 2.8 | 2 | 2.1 | 2684 | 3.7 | 4 | 2.3 | 2612 |
| Iceland | 4.0 | 4 | 2.3 | 3150 | 3.7 | 4 | 2.3 | 1648 | 4.3 | 5 | 2.3 | 1502 |
| Ireland | 3.1 | 3 | 2.2 | 5475 | 2.5 | 2 | 2.0 | 2713 | 3.7 | 4 | 2.2 | 2762 |
| Israel | 2.9 | 2 | 2.3 | 6265 | 2.5 | 2 | 2.3 | 3535 | 3.3 | 3 | 2.2 | 2730 |
| Japan | 2.9 | 2 | 2.7 | 6388 | 2.1 | 1 | 2.5 | 3199 | 3.6 | 4 | 2.8 | 3189 |
| Korea | 2.2 | 2 | 2.2 | 5499 | 1.5 | 1 | 1.8 | 2633 | 3.0 | 2 | 2.2 | 2866 |
| Latvia | 3.1 | 3 | 2.1 | 4667 | 2.8 | 3 | 2.0 | 2345 | 3.5 | 3 | 2.2 | 2322 |
| Lithuania | 3.1 | 3 | 2.2 | 6030 | 2.6 | 2 | 2.0 | 3005 | 3.6 | 3 | 2.2 | 3025 |
| Luxembourg | 2.9 | 3 | 2.2 | 4813 | 2.4 | 2 | 2.0 | 2462 | 3.4 | 3 | 2.3 | 2351 |
| Mexico | 2.9 | 2 | 2.2 | 7006 | 2.4 | 2 | 2.0 | 3550 | 3.3 | 3 | 2.3 | 3456 |
| Montenegro | 3.3 | 3 | 2.4 | 4789 | 2.7 | 2 | 2.2 | 2415 | 3.9 | 4 | 2.4 | 2374 |
| Netherlands | 2.7 | 3 | 1.9 | 5012 | 2.4 | 2 | 1.8 | 2545 | 3.0 | 3 | 1.9 | 2467 |
| New Zealand | 2.9 | 3 | 2.2 | 4100 | 2.6 | 2 | 2.1 | 2062 | 3.2 | 3 | 2.3 | 2038 |
| Norway | 3.4 | 3 | 2.2 | 4982 | 3.1 | 3 | 2.0 | 2512 | 3.6 | 4 | 2.3 | 2470 |
| Peru | 2.6 | 2 | 2.1 | 6343 | 2.0 | 1 | 1.8 | 3099 | 3.1 | 3 | 2.1 | 3244 |
| Poland | 3.6 | 3 | 2.3 | 4318 | 3.2 | 3 | 2.3 | 2148 | 4.0 | 4 | 2.3 | 2170 |
| Portugal | 2.6 | 2 | 2.1 | 6794 | 2.1 | 2 | 2.0 | 3424 | 3.1 | 3 | 2.2 | 3370 |
| Qatar | 2.4 | 2 | 2.2 | 10510 | 1.9 | 1 | 2.0 | 5671 | 3.0 | 3 | 2.3 | 4839 |
| Russian Federation | 3.2 | 3 | 2.2 | 5513 | 2.8 | 3 | 2.1 | 2872 | 3.6 | 3 | 2.2 | 2641 |
| Singapore | 2.2 | 2 | 1.9 | 5968 | 1.8 | 1 | 1.6 | 2909 | 2.6 | 2 | 2.0 | 3059 |
| Slovakia | 3.2 | 3 | 2.3 | 5875 | 2.7 | 2 | 2.1 | 2833 | 3.6 | 3 | 2.3 | 3042 |
| Slovenia | 3.3 | 3 | 2.3 | 6008 | 2.8 | 2 | 2.1 | 2779 | 3.8 | 4 | 2.3 | 3229 |
| Spain | 2.5 | 2 | 2.1 | 6477 | 2.1 | 2 | 1.9 | 3297 | 3.0 | 3 | 2.1 | 3180 |
| Sweden | 3.3 | 3 | 2.2 | 4943 | 3.0 | 3 | 2.0 | 2520 | 3.6 | 3 | 2.3 | 2423 |
| Switzerland | 2.9 | 3 | 2.0 | 5403 | 2.5 | 2 | 1.9 | 2609 | 3.3 | 3 | 2.1 | 2794 |
| Thailand | 2.7 | 2 | 2.1 | 7658 | 2.2 | 2 | 1.8 | 4319 | 3.2 | 3 | 2.3 | 3339 |
| Tunisia | 2.2 | 2 | 2.1 | 4650 | 1.7 | 1 | 1.9 | 2597 | 2.9 | 2 | 2.2 | 2053 |
| Turkey | 2.4 | 2 | 2.2 | 5534 | 1.9 | 1 | 2.0 | 2773 | 2.9 | 2 | 2.2 | 2761 |
| United Arab Emirates | 2.4 | 2 | 2.2 | 13061 | 1.8 | 1 | 2.0 | 6786 | 2.9 | 2 | 2.3 | 6275 |
| United Kingdom | 2.4 | 2 | 2.1 | 12909 | 1.9 | 1 | 1.9 | 6396 | 3.0 | 3 | 2.2 | 6513 |
| United States | 3.5 | 3 | 2.4 | 5294 | 2.9 | 3 | 2.3 | 2668 | 4.1 | 5 | 2.4 | 2626 |
| Uruguay | 2.6 | 2 | 2.2 | 5115 | 2.0 | 2 | 2.0 | 2719 | 3.4 | 3 | 2.3 | 2396 |

Supplementary Table 3. Frequencies of physical activity outcomes by country

| **(a) Physical activity in school, classes per week (overall)** | | | | | | | | |
| --- | --- | --- | --- | --- | --- | --- | --- | --- |
| Country | 0 | 1 | 2 | 3 | 4 | 5 | 6 | 7 |
| Australia | 15.5 | 17.1 | 28.4 | 21.4 | 9.5 | 8.1 | 0.0 | 0.0 |
| Austria | 11.0 | 65.7 | 15.3 | 3.2 | 1.3 | 1.9 | 1.5 | 0.0 |
| Belgium | 1.3 | 52.8 | 38.7 | 2.6 | 1.8 | 2.1 | 0.2 | 0.4 |
| Brazil | 11.5 | 38.6 | 35.2 | 3.4 | 1.7 | 2.9 | 1.3 | 5.4 |
| Bulgaria | 2.1 | 5.5 | 50.7 | 25.6 | 3.8 | 3.9 | 1.9 | 6.4 |
| Canada | 24.3 | 11.5 | 9.4 | 13.8 | 4.4 | 36.7 | 0.0 | 0.0 |
| Chile | 1.0 | 53.9 | 32.4 | 1.9 | 1.8 | 8.8 | 0.0 | 0.0 |
| Chinese Taipei | 0.9 | 22.5 | 71.4 | 2.9 | 0.5 | 1.8 | 0.0 | 0.0 |
| Colombia | 2.9 | 65.3 | 18.6 | 1.5 | 1.2 | 4.5 | 6.0 | 0.0 |
| Costa Rica | 7.7 | 89.2 | 2.0 | 0.2 | 0.1 | 0.8 | 0.0 | 0.0 |
| Croatia | 1.2 | 35.3 | 61.1 | 1.1 | 0.5 | 0.4 | 0.5 | 0.0 |
| Czech Republic | 3.2 | 39.5 | 46.9 | 4.9 | 2.2 | 3.3 | 0.0 | 0.0 |
| Denmark | 2.9 | 72.7 | 13.7 | 4.1 | 2.4 | 4.2 | 0.0 | 0.0 |
| Dominican Republic | 7.8 | 25.4 | 33.0 | 6.2 | 3.0 | 24.6 | 0.0 | 0.0 |
| Estonia | 3.1 | 32.2 | 60.4 | 1.5 | 0.4 | 2.4 | 0.0 | 0.0 |
| Finland | 1.2 | 47.3 | 34.8 | 8.5 | 2.6 | 5.6 | 0.0 | 0.0 |
| France | 3.5 | 70.5 | 17.2 | 3.7 | 1.2 | 3.9 | 0.0 | 0.0 |
| Germany | 1.9 | 66.7 | 25.2 | 3.0 | 1.0 | 1.3 | 1.0 | 0.0 |
| Greece | 3.0 | 6.6 | 72.8 | 5.6 | 1.9 | 2.0 | 1.0 | 7.1 |
| Hong Kong | 1.3 | 89.8 | 7.0 | 0.6 | 0.4 | 0.9 | 0.0 | 0.0 |
| Hungary | 1.2 | 1.7 | 9.3 | 31.3 | 21.6 | 33.3 | 0.5 | 1.2 |
| Iceland | 3.0 | 12.6 | 51.2 | 17.3 | 6.6 | 2.9 | 1.9 | 4.5 |
| Ireland | 9.1 | 76.8 | 10.1 | 1.0 | 0.6 | 2.4 | 0.0 | 0.0 |
| Israel | 9.4 | 26.4 | 47.4 | 3.6 | 2.3 | 2.6 | 1.3 | 6.9 |
| Japan | 0.1 | 5.1 | 40.5 | 51.3 | 2.4 | 0.3 | 0.3 | 0.0 |
| Korea | 0.5 | 7.4 | 73.2 | 13.1 | 3.7 | 2.1 | 0.0 | 0.0 |
| Latvia | 5.3 | 14.3 | 65.4 | 6.9 | 1.8 | 6.3 | 0.0 | 0.0 |
| Lithuania | 7.1 | 5.9 | 75.6 | 4.3 | 1.6 | 5.5 | 0.0 | 0.0 |
| Luxembourg | 2.4 | 58.5 | 27.6 | 5.1 | 2.1 | 4.2 | 0.0 | 0.0 |
| Mexico | 23.1 | 27.1 | 34.3 | 6.0 | 2.1 | 2.4 | 5.0 | 0.0 |
| Montenegro | 3.3 | 5.8 | 53.9 | 8.7 | 3.3 | 4.7 | 2.0 | 18.3 |
| Netherlands | 3.7 | 61.6 | 27.6 | 4.7 | 1.4 | 1.0 | 0.0 | 0.0 |
| New Zealand | 40.3 | 7.0 | 11.9 | 10.0 | 18.6 | 9.7 | 2.4 | 0.0 |
| Norway | 0.7 | 36.3 | 50.2 | 9.8 | 1.3 | 0.9 | 0.3 | 0.5 |
| Peru | 3.0 | 64.3 | 17.1 | 1.7 | 1.1 | 12.7 | 0.0 | 0.0 |
| Poland | 0.6 | 1.0 | 22.8 | 30.5 | 39.8 | 5.4 | 0.0 | 0.0 |
| Portugal | 0.6 | 9.2 | 81.3 | 5.3 | 0.9 | 0.4 | 0.4 | 1.8 |
| Qatar | 12.9 | 42.5 | 17.2 | 8.2 | 4.4 | 14.7 | 0.0 | 0.0 |
| Russian Federation | 3.6 | 3.6 | 17.7 | 62.4 | 3.8 | 1.7 | 1.7 | 5.5 |
| Singapore | 2.4 | 38.9 | 49.4 | 6.9 | 1.0 | 1.4 | 0.0 | 0.0 |
| Slovakia | 4.0 | 14.3 | 65.6 | 10.4 | 2.3 | 3.4 | 0.0 | 0.0 |
| Slovenia | 1.1 | 18.9 | 54.6 | 20.9 | 1.8 | 2.7 | 0.0 | 0.0 |
| Spain | 1.7 | 9.4 | 86.3 | 1.0 | 0.4 | 1.1 | 0.0 | 0.0 |
| Sweden | 2.0 | 16.1 | 64.3 | 5.6 | 2.2 | 1.8 | 0.6 | 7.3 |
| Switzerland | 3.9 | 22.5 | 58.1 | 10.0 | 1.8 | 1.9 | 1.8 | 0.0 |
| Thailand | 8.2 | 74.5 | 9.1 | 2.0 | 0.9 | 5.3 | 0.0 | 0.0 |
| Tunisia | 9.1 | 35.5 | 34.7 | 8.7 | 2.7 | 9.4 | 0.0 | 0.0 |
| Turkey | 5.1 | 46.7 | 23.8 | 2.3 | 4.0 | 18.1 | 0.0 | 0.0 |
| United Arab Emirates | 8.3 | 41.1 | 32.4 | 5.1 | 2.4 | 10.7 | 0.0 | 0.0 |
| United Kingdom | 4.3 | 38.1 | 35.1 | 11.5 | 6.2 | 4.8 | 0.0 | 0.0 |
| United States | 41.3 | 2.6 | 6.3 | 11.6 | 5.1 | 33.1 | 0.0 | 0.0 |
| Uruguay | 19.0 | 11.4 | 53.4 | 4.8 | 1.6 | 3.8 | 6.0 | 0.0 |

| **(b) Physical activity in school, hours per week (overall)** | | | | | | |
| --- | --- | --- | --- | --- | --- | --- |
| Country | 0 hours | 0-1 hours | 1-2 hours | 2-3 hours | 3-4 hours | > 4 hours |
| Australia | 15.4 | 14.0 | 28.0 | 21.1 | 13.0 | 8.5 |
| Austria | 10.9 | 64.9 | 16.2 | 3.4 | 1.5 | 3.1 |
| Belgium | 1.3 | 52.7 | 38.9 | 3.0 | 1.7 | 2.4 |
| Brazil | 11.5 | 38.2 | 36.2 | 3.8 | 2.6 | 7.7 |
| Bulgaria | 1.9 | 5.6 | 66.5 | 13.3 | 5.4 | 7.2 |
| Canada | 24.4 | 2.6 | 12.5 | 9.0 | 11.8 | 39.8 |
| Chile | 1.0 | 37.4 | 39.3 | 10.4 | 6.3 | 5.7 |
| Chinese Taipei | 0.9 | 23.5 | 70.2 | 3.2 | 1.1 | 0.9 |
| Colombia | 2.7 | 55.8 | 28.2 | 2.4 | 2.4 | 8.4 |
| Costa Rica | 7.7 | 89.4 | 2.0 | 0.2 | 0.7 | 0.0 |
| Croatia | 1.2 | 35.2 | 61.0 | 1.7 | 0.6 | 0.4 |
| Czech Republic | 3.2 | 39.7 | 46.9 | 7.0 | 2.9 | 0.4 |
| Denmark | 2.8 | 67.4 | 18.1 | 6.0 | 3.1 | 2.5 |
| Dominican Republic | 8.0 | 26.6 | 32.3 | 9.9 | 9.9 | 13.3 |
| Estonia | 3.1 | 32.0 | 60.5 | 2.1 | 2.0 | 0.2 |
| Finland | 1.1 | 44.7 | 33.6 | 12.8 | 6.2 | 1.5 |
| France | 3.3 | 68.1 | 19.2 | 4.3 | 1.6 | 3.5 |
| Germany | 1.9 | 63.1 | 26.3 | 6.2 | 1.3 | 1.3 |
| Greece | 2.9 | 9.2 | 71.6 | 5.9 | 2.8 | 7.7 |
| Hong Kong | 1.3 | 87.7 | 9.3 | 0.9 | 0.5 | 0.3 |
| Hungary | 1.2 | 1.9 | 10.2 | 51.2 | 33.2 | 2.3 |
| Iceland | 2.9 | 12.5 | 61.6 | 11.5 | 5.6 | 5.8 |
| Ireland | 9.1 | 76.8 | 11.1 | 0.8 | 2.0 | 0.3 |
| Israel | 9.4 | 26.0 | 48.3 | 5.2 | 2.5 | 8.5 |
| Japan | 0.1 | 4.1 | 40.3 | 51.1 | 3.8 | 0.6 |
| Korea | 0.5 | 7.5 | 73.0 | 15.8 | 2.1 | 1.0 |
| Latvia | 5.3 | 14.1 | 71.9 | 2.4 | 5.8 | 0.5 |
| Lithuania | 7.1 | 5.9 | 75.7 | 5.9 | 5.4 | 0.0 |
| Luxembourg | 2.3 | 56.9 | 28.6 | 5.3 | 2.7 | 4.1 |
| Mexico | 23.3 | 27.5 | 33.2 | 6.7 | 3.3 | 6.0 |
| Montenegro | 3.2 | 6.3 | 53.8 | 11.5 | 5.1 | 20.1 |
| Netherlands | 3.7 | 58.3 | 28.1 | 7.5 | 1.4 | 1.1 |
| New Zealand | 40.4 | 6.2 | 11.9 | 10.2 | 19.2 | 12.1 |
| Norway | 0.6 | 34.0 | 48.8 | 13.4 | 1.9 | 1.3 |
| Peru | 3.0 | 61.6 | 20.2 | 3.2 | 6.0 | 5.9 |
| Poland | 0.6 | 1.1 | 23.6 | 68.4 | 6.0 | 0.4 |
| Portugal | 0.6 | 5.9 | 45.6 | 39.5 | 4.8 | 3.7 |
| Qatar | 12.7 | 42.0 | 18.4 | 8.3 | 8.0 | 10.6 |
| Russian Federation | 3.5 | 3.8 | 34.3 | 49.2 | 2.3 | 6.9 |
| Singapore | 2.4 | 48.8 | 39.3 | 6.2 | 1.9 | 1.4 |
| Slovakia | 3.9 | 15.0 | 65.4 | 12.1 | 2.7 | 0.8 |
| Slovenia | 1.1 | 18.8 | 56.0 | 21.4 | 2.5 | 0.2 |
| Spain | 1.7 | 10.3 | 84.6 | 1.8 | 0.6 | 1.0 |
| Sweden | 2.0 | 12.0 | 54.5 | 18.1 | 3.1 | 10.3 |
| Switzerland | 3.9 | 22.2 | 58.3 | 11.6 | 2.1 | 1.9 |
| Thailand | 8.3 | 69.4 | 14.0 | 2.1 | 1.5 | 4.7 |
| Tunisia | 9.1 | 32.6 | 34.0 | 10.6 | 4.9 | 8.8 |
| Turkey | 5.0 | 47.4 | 25.4 | 5.6 | 15.5 | 1.1 |
| United Arab Emirates | 8.2 | 41.1 | 31.4 | 6.8 | 7.4 | 5.1 |
| United Kingdom | 4.2 | 35.7 | 36.4 | 12.8 | 6.6 | 4.4 |
| United States | 41.3 | 2.0 | 7.7 | 7.4 | 15.7 | 25.8 |
| Uruguay | 19.0 | 12.5 | 52.8 | 5.8 | 5.0 | 4.9 |

NB: percent of pupils in each category

| **(c) Out of school moderate activity (60mins+), days/week (overall)** | | | | | | | | |
| --- | --- | --- | --- | --- | --- | --- | --- | --- |
| Country | 0 | 1 | 2 | 3 | 4 | 5 | 6 | 7 |
| Australia | 11.6 | 11.5 | 13.9 | 14.4 | 10.1 | 14.5 | 5.5 | 18.5 |
| Austria | 10.0 | 11.7 | 10.9 | 9.7 | 7.0 | 11.7 | 4.7 | 34.4 |
| Belgium | 13.0 | 15.5 | 12.6 | 10.4 | 6.9 | 13.4 | 4.8 | 23.4 |
| Brazil | 22.8 | 17.5 | 15.9 | 11.4 | 5.9 | 9.0 | 2.7 | 14.8 |
| Bulgaria | 10.9 | 12.1 | 17.1 | 14.4 | 9.2 | 10.1 | 4.8 | 21.4 |
| Canada | 7.1 | 7.0 | 10.7 | 14.0 | 10.8 | 16.6 | 6.2 | 27.8 |
| Chile | 12.3 | 15.3 | 15.2 | 14.1 | 7.3 | 11.6 | 3.5 | 20.5 |
| Chinese Taipei | 15.8 | 11.2 | 13.3 | 8.7 | 3.8 | 16.4 | 3.8 | 27.0 |
| Colombia | 23.9 | 20.7 | 14.0 | 10.1 | 5.0 | 9.6 | 2.6 | 13.9 |
| Costa Rica | 14.7 | 20.2 | 17.3 | 13.3 | 7.3 | 10.2 | 3.0 | 14.0 |
| Croatia | 12.1 | 13.4 | 14.4 | 11.4 | 7.2 | 11.2 | 4.7 | 25.7 |
| Czech Republic | 6.7 | 12.1 | 12.0 | 12.6 | 8.7 | 10.6 | 4.7 | 32.5 |
| Denmark | 6.5 | 7.8 | 9.0 | 9.9 | 8.3 | 17.3 | 8.2 | 33.1 |
| Dominican Republic | 14.0 | 17.2 | 17.8 | 10.6 | 7.1 | 13.1 | 3.5 | 16.7 |
| Estonia | 9.7 | 9.3 | 15.6 | 15.3 | 10.9 | 13.1 | 5.4 | 20.7 |
| Finland | 5.5 | 8.7 | 11.1 | 13.4 | 11.5 | 15.7 | 9.1 | 25.0 |
| France | 13.2 | 14.0 | 13.6 | 10.8 | 8.4 | 8.7 | 4.5 | 26.8 |
| Germany | 5.8 | 8.4 | 10.2 | 10.4 | 8.3 | 13.6 | 6.1 | 37.2 |
| Greece | 13.8 | 13.4 | 16.3 | 15.3 | 9.4 | 9.6 | 4.2 | 18.1 |
| Hong Kong | 17.7 | 15.0 | 12.8 | 10.7 | 5.1 | 12.0 | 3.6 | 23.1 |
| Hungary | 8.6 | 8.7 | 11.8 | 12.5 | 7.6 | 14.5 | 5.1 | 31.3 |
| Iceland | 8.6 | 8.8 | 10.3 | 11.0 | 10.7 | 13.1 | 10.7 | 26.7 |
| Ireland | 9.7 | 13.7 | 16.0 | 14.2 | 9.8 | 12.5 | 6.6 | 17.5 |
| Israel | 19.3 | 13.8 | 13.6 | 11.9 | 7.7 | 6.6 | 14.3 | 12.7 |
| Japan | 27.1 | 6.9 | 5.6 | 5.6 | 3.2 | 14.8 | 10.6 | 26.2 |
| Korea | 19.8 | 12.3 | 13.6 | 10.6 | 5.1 | 14.6 | 4.0 | 20.1 |
| Latvia | 7.3 | 9.0 | 12.5 | 13.8 | 9.9 | 12.6 | 6.0 | 28.8 |
| Lithuania | 8.9 | 10.0 | 12.2 | 12.9 | 8.6 | 12.6 | 4.9 | 29.8 |
| Luxembourg | 12.5 | 15.8 | 15.0 | 12.0 | 9.4 | 9.7 | 4.6 | 20.9 |
| Mexico | 9.9 | 16.3 | 17.2 | 14.6 | 8.0 | 12.9 | 3.9 | 17.2 |
| Montenegro | 5.9 | 9.2 | 14.2 | 14.9 | 10.6 | 12.7 | 5.1 | 27.4 |
| Netherlands | 6.0 | 7.0 | 8.4 | 8.0 | 5.3 | 24.8 | 11.6 | 28.8 |
| New Zealand | 10.6 | 10.3 | 12.3 | 12.6 | 10.1 | 15.6 | 7.0 | 21.3 |
| Norway | 7.1 | 7.3 | 9.2 | 9.8 | 7.7 | 16.6 | 7.7 | 34.6 |
| Peru | 8.8 | 21.0 | 17.5 | 14.0 | 7.3 | 10.8 | 2.9 | 17.7 |
| Poland | 7.0 | 8.0 | 9.4 | 10.6 | 9.3 | 11.2 | 6.9 | 37.7 |
| Portugal | 15.3 | 12.9 | 17.1 | 12.1 | 6.8 | 9.8 | 3.1 | 23.0 |
| Qatar | 23.2 | 16.8 | 14.7 | 13.1 | 8.1 | 7.9 | 2.8 | 13.4 |
| Russian Federation | 6.4 | 8.4 | 12.8 | 16.7 | 10.3 | 8.7 | 7.6 | 29.0 |
| Singapore | 15.2 | 14.9 | 14.4 | 10.0 | 5.0 | 11.5 | 3.1 | 25.8 |
| Slovakia | 8.2 | 11.0 | 13.0 | 11.7 | 8.9 | 12.2 | 4.2 | 30.9 |
| Slovenia | 6.3 | 12.7 | 14.3 | 14.7 | 9.3 | 12.6 | 5.7 | 24.3 |
| Spain | 16.6 | 11.7 | 16.8 | 13.4 | 8.9 | 10.5 | 3.9 | 18.3 |
| Sweden | 8.7 | 9.3 | 10.3 | 10.9 | 8.9 | 15.0 | 7.2 | 29.7 |
| Switzerland | 7.5 | 11.5 | 11.5 | 10.5 | 7.2 | 12.9 | 6.6 | 32.4 |
| Thailand | 4.7 | 15.8 | 16.2 | 15.6 | 6.4 | 15.0 | 1.8 | 24.5 |
| Tunisia | 20.6 | 21.0 | 17.8 | 13.6 | 7.2 | 4.8 | 3.9 | 11.0 |
| Turkey | 17.7 | 19.7 | 15.5 | 10.8 | 6.1 | 8.9 | 1.6 | 19.6 |
| United Arab Emirates | 27.3 | 16.8 | 14.5 | 10.9 | 6.9 | 7.7 | 2.5 | 13.5 |
| United Kingdom | 11.4 | 14.3 | 12.9 | 11.4 | 8.0 | 13.5 | 4.9 | 23.5 |
| United States | 10.7 | 7.3 | 10.1 | 11.4 | 8.4 | 16.5 | 7.0 | 28.5 |
| Uruguay | 15.9 | 12.3 | 16.2 | 12.1 | 7.6 | 12.4 | 5.7 | 17.8 |

NB: percent of pupils in each category

| **(d) Out of school vigorous activity (20mins+), days/week (overall)** | | | | | | | | |
| --- | --- | --- | --- | --- | --- | --- | --- | --- |
| Country | 0 | 1 | 2 | 3 | 4 | 5 | 6 | 7 |
| Australia | 16.9 | 14.0 | 16.9 | 16.0 | 11.7 | 9.8 | 4.7 | 10.1 |
| Austria | 20.8 | 19.1 | 17.9 | 14.8 | 10.2 | 7.1 | 3.4 | 6.5 |
| Belgium | 18.9 | 19.1 | 17.5 | 15.8 | 10.1 | 8.1 | 3.6 | 6.9 |
| Brazil | 35.1 | 16.2 | 13.5 | 9.9 | 5.7 | 7.8 | 2.9 | 8.9 |
| Bulgaria | 17.4 | 16.1 | 17.1 | 14.6 | 8.6 | 9.3 | 4.8 | 12.1 |
| Canada | 15.2 | 11.8 | 14.6 | 14.6 | 11.5 | 13.3 | 5.8 | 13.2 |
| Chile | 21.5 | 19.8 | 17.8 | 14.3 | 8.6 | 6.5 | 3.0 | 8.5 |
| Chinese Taipei | 18.4 | 15.6 | 25.3 | 12.3 | 6.3 | 8.5 | 2.9 | 10.8 |
| Colombia | 19.4 | 24.2 | 16.9 | 11.9 | 7.6 | 8.2 | 3.5 | 8.3 |
| Costa Rica | 27.9 | 20.1 | 15.3 | 13.3 | 6.8 | 6.2 | 2.9 | 7.5 |
| Croatia | 19.6 | 15.9 | 16.9 | 13.2 | 8.8 | 8.9 | 4.7 | 11.9 |
| Czech Republic | 11.5 | 16.0 | 16.2 | 17.0 | 12.5 | 10.0 | 5.5 | 11.4 |
| Denmark | 11.1 | 11.5 | 14.8 | 17.5 | 13.1 | 12.5 | 6.3 | 13.1 |
| Dominican Republic | 16.8 | 17.8 | 17.1 | 12.7 | 8.7 | 10.7 | 4.5 | 11.8 |
| Estonia | 13.1 | 13.2 | 18.1 | 16.9 | 12.4 | 11.3 | 4.8 | 10.3 |
| Finland | 12.1 | 15.2 | 18.4 | 17.0 | 11.8 | 11.3 | 7.4 | 6.8 |
| France | 23.2 | 21.2 | 19.1 | 14.3 | 8.4 | 5.1 | 2.6 | 6.0 |
| Germany | 11.8 | 14.2 | 20.2 | 19.1 | 13.6 | 9.7 | 3.9 | 7.7 |
| Greece | 18.9 | 14.6 | 16.4 | 15.3 | 10.2 | 9.6 | 4.8 | 10.2 |
| Hong Kong | 22.7 | 23.2 | 17.3 | 13.0 | 6.1 | 7.1 | 2.4 | 8.2 |
| Hungary | 13.8 | 11.6 | 16.8 | 16.6 | 11.1 | 11.6 | 5.1 | 13.5 |
| Iceland | 9.7 | 8.6 | 11.3 | 11.8 | 12.1 | 14.8 | 11.4 | 20.3 |
| Ireland | 14.3 | 13.0 | 16.1 | 15.7 | 12.6 | 11.3 | 6.4 | 10.5 |
| Israel | 20.1 | 15.0 | 15.3 | 12.5 | 9.0 | 6.7 | 14.0 | 7.4 |
| Japan | 33.6 | 11.0 | 8.8 | 8.1 | 3.7 | 6.8 | 10.8 | 17.1 |
| Korea | 27.8 | 16.2 | 20.0 | 12.5 | 5.8 | 7.2 | 2.5 | 8.0 |
| Latvia | 12.5 | 13.1 | 17.1 | 17.2 | 11.9 | 12.3 | 5.3 | 10.6 |
| Lithuania | 14.2 | 12.7 | 17.7 | 16.5 | 11.3 | 11.7 | 5.0 | 11.0 |
| Luxembourg | 16.6 | 15.8 | 16.9 | 15.6 | 11.4 | 8.7 | 4.7 | 10.4 |
| Mexico | 14.8 | 18.3 | 18.9 | 13.5 | 8.4 | 11.5 | 4.5 | 10.1 |
| Montenegro | 14.7 | 13.6 | 15.2 | 14.2 | 9.0 | 10.5 | 5.4 | 17.5 |
| Netherlands | 16.1 | 14.7 | 16.2 | 23.0 | 13.3 | 8.0 | 4.3 | 4.4 |
| New Zealand | 19.3 | 11.8 | 16.8 | 14.4 | 12.1 | 10.6 | 6.0 | 9.0 |
| Norway | 10.9 | 11.9 | 15.0 | 17.0 | 13.4 | 12.8 | 7.0 | 12.0 |
| Peru | 14.6 | 23.9 | 19.7 | 14.5 | 8.2 | 7.2 | 3.3 | 8.6 |
| Poland | 11.4 | 11.6 | 13.1 | 15.5 | 11.8 | 10.7 | 6.6 | 19.2 |
| Portugal | 21.2 | 14.8 | 19.7 | 14.1 | 10.7 | 7.7 | 3.3 | 8.5 |
| Qatar | 25.4 | 18.0 | 16.0 | 12.1 | 8.1 | 7.6 | 3.6 | 9.1 |
| Russian Federation | 12.3 | 12.7 | 15.3 | 21.4 | 11.2 | 8.7 | 6.0 | 12.5 |
| Singapore | 18.7 | 23.5 | 22.1 | 14.9 | 7.8 | 6.1 | 2.1 | 4.9 |
| Slovakia | 12.5 | 15.3 | 16.8 | 16.4 | 10.2 | 9.7 | 4.8 | 14.3 |
| Slovenia | 11.3 | 14.6 | 15.8 | 15.1 | 11.5 | 10.7 | 6.6 | 14.4 |
| Spain | 21.7 | 13.1 | 19.8 | 14.9 | 12.7 | 7.7 | 3.8 | 6.3 |
| Sweden | 12.4 | 12.5 | 15.2 | 15.3 | 13.5 | 12.2 | 7.5 | 11.4 |
| Switzerland | 12.2 | 15.6 | 18.5 | 18.6 | 12.7 | 10.3 | 4.6 | 7.5 |
| Thailand | 11.6 | 24.0 | 21.0 | 15.8 | 6.3 | 9.4 | 1.7 | 10.1 |
| Tunisia | 25.6 | 20.5 | 17.2 | 12.8 | 8.0 | 5.0 | 3.1 | 7.8 |
| Turkey | 21.8 | 21.3 | 18.8 | 13.1 | 6.7 | 6.4 | 2.2 | 9.7 |
| United Arab Emirates | 26.2 | 19.7 | 15.6 | 11.4 | 7.3 | 7.5 | 2.9 | 9.5 |
| United Kingdom | 21.5 | 19.5 | 17.6 | 14.2 | 9.0 | 7.9 | 3.6 | 6.7 |
| United States | 16.7 | 9.9 | 11.9 | 12.3 | 8.7 | 16.4 | 7.6 | 16.6 |
| Uruguay | 24.4 | 12.4 | 17.3 | 14.7 | 7.9 | 9.2 | 5.8 | 8.4 |

NB: percent of pupils in each category

Supplementary Table 4. Tabulations of physical activity outcomes by gender.

| **(a) Physical activity in school, classes per week** | | | | | | | | | | | | | | | |  |  |
| --- | --- | --- | --- | --- | --- | --- | --- | --- | --- | --- | --- | --- | --- | --- | --- | --- | --- |
|  | Female | |  |  |  |  |  |  |  | Male |  |  |  |  |  |  |  |
| Country | 0 | 1 | 2 | 3 | 4 | 5 | 6 | 7 |  | 0 | 1 | 2 | 3 | 4 | 5 | 6 | 7 |
| Australia | 17.8 | 18.7 | 28.9 | 20.8 | 8.0 | 5.8 | 0.0 | 0.0 |  | 15.6 | 27.8 | 22.1 | 11.0 | 10.4 | 0.0 | 0.0 | 15.6 |
| Austria | 9.3 | 68.1 | 16.7 | 3.1 | 1.1 | 0.8 | 0.9 | 0.0 |  | 63.3 | 13.9 | 3.3 | 1.5 | 3.0 | 2.2 | 0.0 | 63.3 |
| Belgium | 1.4 | 55.6 | 38.0 | 2.0 | 1.0 | 1.4 | 0.2 | 0.4 |  | 50.0 | 39.4 | 3.2 | 2.7 | 2.9 | 0.1 | 0.5 | 50.0 |
| Brazil | 12.4 | 39.2 | 35.4 | 3.3 | 1.6 | 2.4 | 1.2 | 4.5 |  | 37.9 | 35.0 | 3.6 | 1.7 | 3.4 | 1.5 | 6.5 | 37.9 |
| Bulgaria | 2.0 | 4.6 | 55.7 | 25.6 | 3.4 | 2.5 | 1.2 | 4.9 |  | 6.4 | 46.1 | 25.6 | 4.1 | 5.2 | 2.6 | 7.8 | 6.4 |
| Canada | 28.7 | 12.8 | 9.2 | 13.8 | 4.3 | 31.3 | 0.0 | 0.0 |  | 10.2 | 9.6 | 13.9 | 4.4 | 42.2 | 0.0 | 0.0 | 10.2 |
| Chile | 1.4 | 55.0 | 31.7 | 2.1 | 2.1 | 7.8 | 0.0 | 0.0 |  | 52.9 | 33.2 | 1.8 | 1.5 | 9.9 | 0.0 | 0.0 | 52.9 |
| Chinese Taipei | 0.7 | 23.6 | 71.6 | 2.4 | 0.3 | 1.4 | 0.0 | 0.0 |  | 21.4 | 71.2 | 3.4 | 0.6 | 2.2 | 0.0 | 0.0 | 21.4 |
| Colombia | 3.1 | 67.0 | 18.9 | 1.4 | 0.9 | 3.6 | 5.2 | 0.0 |  | 63.5 | 18.3 | 1.5 | 1.5 | 5.6 | 6.9 | 0.0 | 63.5 |
| Costa Rica | 7.2 | 90.0 | 2.2 | 0.1 | 0.0 | 0.5 | 0.0 | 0.0 |  | 88.3 | 1.8 | 0.3 | 0.2 | 1.2 | 0.0 | 0.0 | 88.3 |
| Croatia | 1.1 | 34.7 | 62.5 | 0.9 | 0.3 | 0.2 | 0.3 | 0.0 |  | 36.0 | 59.6 | 1.3 | 0.7 | 0.5 | 0.6 | 0.0 | 36.0 |
| Czech Republic | 3.9 | 42.9 | 45.8 | 4.2 | 1.3 | 1.9 | 0.0 | 0.0 |  | 36.3 | 47.9 | 5.6 | 3.0 | 4.6 | 0.0 | 0.0 | 36.3 |
| Denmark | 2.8 | 74.3 | 12.7 | 3.7 | 2.3 | 4.2 | 0.0 | 0.0 |  | 71.2 | 14.7 | 4.5 | 2.5 | 4.2 | 0.0 | 0.0 | 71.2 |
| Dominican Republic | 8.1 | 25.4 | 36.1 | 6.2 | 2.9 | 21.2 | 0.0 | 0.0 |  | 25.4 | 29.7 | 6.2 | 3.1 | 28.1 | 0.0 | 0.0 | 25.4 |
| Estonia | 2.9 | 33.5 | 61.1 | 1.3 | 0.3 | 1.0 | 0.0 | 0.0 |  | 30.9 | 59.8 | 1.7 | 0.6 | 3.8 | 0.0 | 0.0 | 30.9 |
| Finland | 1.2 | 56.6 | 30.6 | 5.9 | 2.0 | 3.7 | 0.0 | 0.0 |  | 38.5 | 38.8 | 10.9 | 3.2 | 7.4 | 0.0 | 0.0 | 38.5 |
| France | 3.9 | 72.6 | 16.2 | 3.6 | 0.5 | 3.3 | 0.0 | 0.0 |  | 68.2 | 18.2 | 3.8 | 2.0 | 4.6 | 0.0 | 0.0 | 68.2 |
| Germany | 2.0 | 68.5 | 24.0 | 2.7 | 0.9 | 1.3 | 0.6 | 0.0 |  | 64.7 | 26.5 | 3.3 | 1.1 | 1.3 | 1.3 | 0.0 | 64.7 |
| Greece | 3.6 | 7.0 | 71.5 | 5.3 | 1.7 | 2.2 | 1.1 | 7.6 |  | 6.3 | 73.9 | 5.9 | 2.2 | 1.8 | 0.9 | 6.7 | 6.3 |
| Hong Kong | 1.3 | 91.0 | 6.4 | 0.3 | 0.4 | 0.6 | 0.0 | 0.0 |  | 88.6 | 7.6 | 0.8 | 0.3 | 1.3 | 0.0 | 0.0 | 88.6 |
| Hungary | 1.4 | 1.4 | 8.0 | 30.8 | 23.2 | 34.0 | 0.4 | 0.8 |  | 1.9 | 10.5 | 31.8 | 20.0 | 32.6 | 0.5 | 1.7 | 1.9 |
| Iceland | 3.8 | 13.8 | 50.8 | 16.9 | 6.5 | 2.5 | 1.9 | 3.7 |  | 11.3 | 51.7 | 17.8 | 6.7 | 3.4 | 1.8 | 5.3 | 11.3 |
| Ireland | 8.7 | 77.6 | 10.7 | 0.8 | 0.4 | 1.8 | 0.0 | 0.0 |  | 76.0 | 9.6 | 1.2 | 0.7 | 3.0 | 0.0 | 0.0 | 76.0 |
| Israel | 7.4 | 30.5 | 49.0 | 3.1 | 1.7 | 1.7 | 1.2 | 5.4 |  | 22.1 | 45.7 | 4.3 | 2.9 | 3.6 | 1.5 | 8.4 | 22.1 |
| Japan | 0.1 | 4.5 | 43.5 | 49.9 | 1.8 | 0.3 | 0.1 | 0.0 |  | 5.7 | 37.7 | 52.7 | 3.0 | 0.4 | 0.5 | 0.0 | 5.7 |
| Korea | 0.1 | 7.8 | 75.7 | 11.9 | 3.2 | 1.2 | 0.0 | 0.0 |  | 6.9 | 70.8 | 14.2 | 4.2 | 2.9 | 0.0 | 0.0 | 6.9 |
| Latvia | 7.0 | 17.0 | 65.6 | 6.1 | 1.3 | 3.0 | 0.0 | 0.0 |  | 11.6 | 65.1 | 7.7 | 2.3 | 9.6 | 0.0 | 0.0 | 11.6 |
| Lithuania | 7.3 | 6.8 | 79.4 | 3.4 | 0.9 | 2.3 | 0.0 | 0.0 |  | 5.1 | 71.7 | 5.2 | 2.4 | 8.7 | 0.0 | 0.0 | 5.1 |
| Luxembourg | 1.8 | 62.9 | 26.2 | 5.0 | 1.4 | 2.7 | 0.0 | 0.0 |  | 54.0 | 29.0 | 5.2 | 2.9 | 5.8 | 0.0 | 0.0 | 54.0 |
| Mexico | 23.4 | 27.9 | 33.6 | 6.2 | 2.0 | 2.1 | 4.7 | 0.0 |  | 26.2 | 35.0 | 5.7 | 2.2 | 2.7 | 5.3 | 0.0 | 26.2 |
| Montenegro | 3.1 | 5.4 | 56.7 | 8.5 | 2.7 | 3.8 | 2.3 | 17.5 |  | 6.2 | 50.9 | 8.9 | 3.9 | 5.6 | 1.8 | 19.1 | 6.2 |
| Netherlands | 2.9 | 64.6 | 26.8 | 3.8 | 1.3 | 0.6 | 0.0 | 0.0 |  | 58.5 | 28.4 | 5.6 | 1.5 | 1.4 | 0.0 | 0.0 | 58.5 |
| New Zealand | 45.8 | 6.6 | 12.3 | 10.3 | 16.7 | 6.9 | 1.4 | 0.0 |  | 7.5 | 11.6 | 9.6 | 20.5 | 12.5 | 3.4 | 0.0 | 7.5 |
| Norway | 0.6 | 38.1 | 51.6 | 7.5 | 1.2 | 0.6 | 0.1 | 0.2 |  | 34.4 | 48.8 | 12.2 | 1.5 | 1.3 | 0.5 | 0.7 | 34.4 |
| Peru | 3.1 | 65.6 | 18.7 | 1.4 | 1.2 | 10.1 | 0.0 | 0.0 |  | 63.1 | 15.6 | 2.1 | 1.1 | 15.2 | 0.0 | 0.0 | 63.1 |
| Poland | 0.7 | 1.1 | 23.3 | 29.0 | 41.8 | 4.1 | 0.0 | 0.0 |  | 0.8 | 22.3 | 31.9 | 37.8 | 6.7 | 0.0 | 0.0 | 0.8 |
| Portugal | 0.7 | 8.1 | 82.7 | 5.4 | 0.6 | 0.4 | 0.3 | 1.8 |  | 10.3 | 79.8 | 5.2 | 1.3 | 0.5 | 0.4 | 1.9 | 10.3 |
| Qatar | 14.4 | 43.4 | 17.1 | 8.2 | 4.3 | 12.6 | 0.0 | 0.0 |  | 41.6 | 17.3 | 8.3 | 4.5 | 17.1 | 0.0 | 0.0 | 41.6 |
| Russian Federation | 4.1 | 4.6 | 18.4 | 63.0 | 3.5 | 1.2 | 1.5 | 3.8 |  | 2.6 | 17.0 | 61.7 | 4.2 | 2.2 | 2.0 | 7.2 | 2.6 |
| Singapore | 3.1 | 37.7 | 49.3 | 8.3 | 1.0 | 0.6 | 0.0 | 0.0 |  | 40.1 | 49.5 | 5.7 | 1.0 | 2.0 | 0.0 | 0.0 | 40.1 |
| Slovakia | 5.1 | 16.1 | 65.8 | 10.1 | 1.1 | 1.8 | 0.0 | 0.0 |  | 12.7 | 65.5 | 10.7 | 3.3 | 4.8 | 0.0 | 0.0 | 12.7 |
| Slovenia | 0.7 | 15.3 | 58.0 | 22.9 | 1.6 | 1.5 | 0.0 | 0.0 |  | 22.4 | 51.4 | 19.1 | 1.9 | 3.7 | 0.0 | 0.0 | 22.4 |
| Spain | 1.6 | 10.4 | 85.7 | 1.2 | 0.4 | 0.7 | 0.0 | 0.0 |  | 8.4 | 86.9 | 0.8 | 0.4 | 1.6 | 0.0 | 0.0 | 8.4 |
| Sweden | 2.7 | 18.2 | 66.1 | 4.5 | 2.1 | 1.6 | 0.8 | 4.0 |  | 14.0 | 62.4 | 6.7 | 2.4 | 2.0 | 0.5 | 10.6 | 14.0 |
| Switzerland | 3.3 | 21.8 | 60.5 | 10.1 | 1.6 | 1.5 | 1.1 | 0.0 |  | 23.1 | 55.9 | 9.8 | 1.9 | 2.3 | 2.5 | 0.0 | 23.1 |
| Thailand | 7.4 | 77.9 | 8.7 | 1.7 | 0.6 | 3.7 | 0.0 | 0.0 |  | 70.1 | 9.6 | 2.3 | 1.2 | 7.5 | 0.0 | 0.0 | 70.1 |
| United Arab Emirates | 10.2 | 37.6 | 35.1 | 7.7 | 2.2 | 7.1 | 0.0 | 0.0 |  | 32.9 | 34.2 | 9.8 | 3.3 | 12.0 | 0.0 | 0.0 | 32.9 |
| Tunisia | 7.4 | 45.1 | 23.5 | 2.6 | 4.8 | 16.6 | 0.0 | 0.0 |  | 48.4 | 24.0 | 2.0 | 3.3 | 19.5 | 0.0 | 0.0 | 48.4 |
| Turkey | 10.0 | 41.9 | 34.9 | 4.3 | 2.1 | 6.9 | 0.0 | 0.0 |  | 40.1 | 29.8 | 6.0 | 2.7 | 15.0 | 0.0 | 0.0 | 40.1 |
| United Kingdom | 5.0 | 43.0 | 34.7 | 9.2 | 4.5 | 3.6 | 0.0 | 0.0 |  | 33.3 | 35.4 | 13.8 | 7.9 | 6.1 | 0.0 | 0.0 | 33.3 |
| United States | 46.4 | 2.7 | 6.5 | 11.6 | 5.2 | 27.6 | 0.0 | 0.0 |  | 2.5 | 6.0 | 11.7 | 5.0 | 38.8 | 0.0 | 0.0 | 2.5 |
| Uruguay | 20.0 | 11.2 | 56.5 | 3.7 | 1.4 | 2.8 | 4.4 | 0.0 |  | 11.6 | 49.7 | 6.0 | 2.0 | 5.1 | 7.8 | 0.0 | 11.6 |

NB: percent of pupils in each category

| **(b) Physical activity in school, hours per week** | | | | | | | | | | | | | |
| --- | --- | --- | --- | --- | --- | --- | --- | --- | --- | --- | --- | --- | --- |
|  | Female |  |  |  |  |  |  | Male |  |  |  |  |  |
| Country | 0 hours | 0-1 hours | 1-2 hours | 2-3 hours | 3-4 hours | > 4 hours |  | 0 hours | 0-1 hours | 1-2 hours | 2-3 hours | 3-4 hours | > 4 hours |
| Australia | 17.7 | 15.3 | 28.3 | 21.0 | 11.5 | 6.2 |  | 13.1 | 12.7 | 27.7 | 21.2 | 14.5 | 10.8 |
| Austria | 9.1 | 67.4 | 17.5 | 3.5 | 1.1 | 1.4 |  | 12.6 | 62.4 | 14.9 | 3.4 | 1.9 | 4.8 |
| Belgium | 1.3 | 55.6 | 38.1 | 2.2 | 0.9 | 1.8 |  | 1.3 | 49.7 | 39.7 | 3.7 | 2.5 | 3.1 |
| Brazil | 12.5 | 39.2 | 36.3 | 3.5 | 2.2 | 6.2 |  | 10.4 | 37.2 | 36.0 | 4.1 | 2.9 | 9.4 |
| Bulgaria | 1.9 | 4.5 | 73.5 | 11.2 | 3.5 | 5.4 |  | 2.0 | 6.7 | 60.1 | 15.2 | 7.1 | 8.9 |
| Canada | 28.7 | 2.6 | 13.5 | 8.9 | 11.7 | 34.6 |  | 19.9 | 2.5 | 11.4 | 9.0 | 11.9 | 45.3 |
| Chile | 1.3 | 40.6 | 37.6 | 9.9 | 5.8 | 4.8 |  | 0.7 | 34.2 | 40.9 | 10.8 | 6.9 | 6.5 |
| Chinese Taipei | 0.7 | 24.7 | 70.4 | 2.6 | 0.8 | 0.8 |  | 1.2 | 22.4 | 70.1 | 3.8 | 1.4 | 1.1 |
| Colombia | 2.9 | 59.6 | 27.0 | 2.3 | 2.0 | 6.3 |  | 2.5 | 51.5 | 29.7 | 2.5 | 2.9 | 10.8 |
| Costa Rica | 7.2 | 90.1 | 2.2 | 0.2 | 0.3 | 0.0 |  | 8.2 | 88.6 | 1.8 | 0.3 | 1.1 | 0.0 |
| Croatia | 1.1 | 34.6 | 62.4 | 1.3 | 0.3 | 0.3 |  | 1.3 | 35.8 | 59.5 | 2.1 | 0.8 | 0.5 |
| Czech Republic | 3.9 | 43.2 | 45.6 | 5.5 | 1.6 | 0.2 |  | 2.4 | 36.3 | 48.2 | 8.4 | 4.2 | 0.5 |
| Denmark | 2.8 | 68.8 | 17.5 | 5.4 | 2.9 | 2.5 |  | 2.9 | 66.0 | 18.7 | 6.6 | 3.3 | 2.6 |
| Dominican Republic | 8.6 | 27.2 | 34.5 | 10.6 | 9.6 | 9.5 |  | 7.4 | 25.9 | 30.0 | 9.1 | 10.3 | 17.3 |
| Estonia | 2.9 | 33.2 | 61.3 | 1.7 | 0.8 | 0.1 |  | 3.2 | 30.9 | 59.8 | 2.6 | 3.2 | 0.4 |
| Finland | 1.2 | 53.5 | 29.5 | 10.5 | 4.5 | 0.8 |  | 1.1 | 36.2 | 37.6 | 15.1 | 7.9 | 2.1 |
| France | 3.7 | 70.7 | 17.5 | 4.5 | 0.8 | 2.7 |  | 2.8 | 65.2 | 21.0 | 4.2 | 2.5 | 4.3 |
| Germany | 2.0 | 64.3 | 25.6 | 6.0 | 1.0 | 1.1 |  | 1.8 | 61.8 | 27.1 | 6.5 | 1.5 | 1.4 |
| Greece | 3.4 | 9.2 | 70.6 | 5.5 | 3.1 | 8.2 |  | 2.3 | 9.1 | 72.6 | 6.3 | 2.5 | 7.2 |
| Hong Kong | 1.3 | 88.8 | 8.8 | 0.6 | 0.5 | 0.0 |  | 1.4 | 86.7 | 9.8 | 1.2 | 0.5 | 0.5 |
| Hungary | 1.3 | 2.0 | 8.8 | 52.4 | 33.9 | 1.5 |  | 1.0 | 1.9 | 11.6 | 49.9 | 32.4 | 3.2 |
| Iceland | 3.9 | 13.6 | 61.3 | 11.0 | 5.3 | 4.9 |  | 1.9 | 11.3 | 62.0 | 12.2 | 5.9 | 6.7 |
| Ireland | 8.7 | 77.6 | 11.3 | 0.6 | 1.6 | 0.1 |  | 9.5 | 76.0 | 10.8 | 1.0 | 2.4 | 0.4 |
| Israel | 7.4 | 30.1 | 49.9 | 4.3 | 1.9 | 6.5 |  | 11.6 | 21.7 | 46.5 | 6.3 | 3.2 | 10.7 |
| Japan | 0.1 | 3.6 | 43.1 | 49.5 | 3.3 | 0.4 |  | 0.1 | 4.6 | 37.4 | 52.6 | 4.4 | 0.9 |
| Korea | 0.1 | 8.1 | 75.6 | 14.1 | 1.5 | 0.5 |  | 0.9 | 7.0 | 70.6 | 17.3 | 2.7 | 1.5 |
| Latvia | 7.0 | 16.9 | 71.4 | 1.6 | 3.0 | 0.1 |  | 3.5 | 11.2 | 72.4 | 3.2 | 8.7 | 0.8 |
| Lithuania | 7.3 | 6.8 | 79.5 | 4.2 | 2.2 | 0.0 |  | 6.9 | 5.0 | 71.9 | 7.6 | 8.6 | 0.0 |
| Luxembourg | 1.8 | 61.5 | 27.2 | 5.0 | 1.8 | 2.8 |  | 2.9 | 52.2 | 30.1 | 5.6 | 3.6 | 5.5 |
| Mexico | 23.5 | 29.3 | 31.9 | 6.9 | 3.0 | 5.3 |  | 23.0 | 25.7 | 34.4 | 6.5 | 3.6 | 6.7 |
| Montenegro | 3.1 | 5.7 | 56.9 | 10.6 | 4.1 | 19.6 |  | 3.4 | 6.9 | 50.6 | 12.3 | 6.1 | 20.7 |
| Netherlands | 2.9 | 60.9 | 27.3 | 6.9 | 1.2 | 0.9 |  | 4.5 | 55.5 | 28.8 | 8.2 | 1.6 | 1.3 |
| New Zealand | 45.6 | 5.7 | 12.2 | 10.8 | 16.6 | 9.0 |  | 35.1 | 6.7 | 11.6 | 9.6 | 21.8 | 15.2 |
| Norway | 0.6 | 36.1 | 50.6 | 10.9 | 1.2 | 0.8 |  | 0.7 | 32.0 | 47.0 | 15.9 | 2.5 | 1.9 |
| Peru | 3.1 | 63.5 | 21.5 | 2.7 | 4.5 | 4.8 |  | 2.9 | 59.8 | 19.1 | 3.8 | 7.5 | 6.9 |
| Poland | 0.7 | 1.2 | 23.8 | 69.5 | 4.7 | 0.1 |  | 0.5 | 1.0 | 23.4 | 67.2 | 7.2 | 0.7 |
| Portugal | 0.7 | 5.1 | 46.2 | 40.8 | 3.8 | 3.4 |  | 0.5 | 6.6 | 45.0 | 38.2 | 5.8 | 4.0 |
| Qatar | 14.3 | 43.0 | 18.1 | 8.4 | 7.6 | 8.6 |  | 10.8 | 40.9 | 18.8 | 8.3 | 8.4 | 12.9 |
| Russian Federation | 4.1 | 4.8 | 35.9 | 48.4 | 1.8 | 5.1 |  | 3.0 | 2.7 | 32.5 | 50.1 | 3.0 | 8.8 |
| Singapore | 3.1 | 46.5 | 41.2 | 6.5 | 1.9 | 0.8 |  | 1.8 | 50.9 | 37.5 | 6.0 | 1.9 | 1.9 |
| Slovakia | 5.1 | 16.8 | 65.7 | 10.8 | 1.3 | 0.3 |  | 2.9 | 13.3 | 65.1 | 13.3 | 4.1 | 1.3 |
| Slovenia | 0.7 | 15.2 | 59.0 | 23.5 | 1.6 | 0.0 |  | 1.4 | 22.3 | 53.1 | 19.3 | 3.5 | 0.4 |
| Spain | 1.5 | 11.3 | 84.5 | 1.6 | 0.5 | 0.5 |  | 1.8 | 9.2 | 84.8 | 1.9 | 0.7 | 1.5 |
| Sweden | 2.5 | 14.1 | 56.7 | 17.0 | 2.5 | 7.1 |  | 1.4 | 9.9 | 52.2 | 19.3 | 3.7 | 13.6 |
| Switzerland | 3.3 | 21.6 | 60.7 | 11.7 | 1.6 | 1.1 |  | 4.5 | 22.8 | 56.0 | 11.4 | 2.6 | 2.7 |
| Thailand | 7.5 | 73.2 | 12.9 | 2.1 | 1.3 | 3.1 |  | 9.4 | 64.4 | 15.4 | 2.2 | 1.9 | 6.7 |
| Tunisia | 10.3 | 34.9 | 34.4 | 9.7 | 4.2 | 6.6 |  | 7.6 | 29.9 | 33.7 | 11.7 | 5.6 | 11.5 |
| Turkey | 7.4 | 45.1 | 25.7 | 6.2 | 14.4 | 1.2 |  | 2.7 | 49.7 | 25.1 | 4.9 | 16.6 | 1.0 |
| United Arab Emirates | 9.9 | 41.8 | 33.8 | 5.8 | 5.4 | 3.2 |  | 6.3 | 40.2 | 28.8 | 7.8 | 9.7 | 7.2 |
| United Kingdom | 4.9 | 39.5 | 37.5 | 9.9 | 5.1 | 3.2 |  | 3.4 | 31.8 | 35.2 | 15.8 | 8.2 | 5.6 |
| United States | 46.6 | 2.2 | 7.7 | 8.1 | 13.9 | 21.5 |  | 36.1 | 1.8 | 7.6 | 6.8 | 17.5 | 30.2 |
| Uruguay | 19.9 | 12.6 | 55.9 | 4.9 | 3.4 | 3.4 |  | 18.1 | 12.5 | 49.2 | 6.9 | 6.8 | 6.6 |

NB: percent of pupils in each category

| **(c) Out of school moderate activity (60mins+), days/week** | | | | | | | | | | | | | | | | | |
| --- | --- | --- | --- | --- | --- | --- | --- | --- | --- | --- | --- | --- | --- | --- | --- | --- | --- |
|  | Female | |  |  |  |  |  |  |  | Male |  |  |  |  |  |  |  |
| Country | 0 | 1 | 2 | 3 | 4 | 5 | 6 | 7 |  | 0 | 1 | 2 | 3 | 4 | 5 | 6 | 7 |
| Australia | 13.1 | 12.9 | 15.4 | 15.5 | 10.0 | 14.5 | 5.4 | 13.2 |  | 10.1 | 10.0 | 12.4 | 13.3 | 10.2 | 14.5 | 5.6 | 23.9 |
| Austria | 9.5 | 12.5 | 11.6 | 10.2 | 7.2 | 11.3 | 5.0 | 32.8 |  | 10.4 | 11.0 | 10.1 | 9.1 | 6.8 | 12.2 | 4.4 | 36.0 |
| Belgium | 14.4 | 16.6 | 13.3 | 10.1 | 6.4 | 13.1 | 4.4 | 21.7 |  | 11.6 | 14.5 | 11.9 | 10.7 | 7.4 | 13.7 | 5.2 | 25.0 |
| Brazil | 27.2 | 17.4 | 15.6 | 11.0 | 5.4 | 8.9 | 2.1 | 12.6 |  | 18.0 | 17.6 | 16.2 | 11.9 | 6.6 | 9.1 | 3.4 | 17.3 |
| Bulgaria | 10.8 | 11.6 | 16.1 | 15.1 | 9.8 | 10.1 | 5.3 | 21.2 |  | 10.9 | 12.6 | 18.1 | 13.7 | 8.6 | 10.1 | 4.4 | 21.6 |
| Canada | 7.8 | 8.3 | 12.3 | 15.8 | 11.2 | 16.2 | 5.6 | 22.9 |  | 6.3 | 5.7 | 9.0 | 12.0 | 10.4 | 16.9 | 6.8 | 32.8 |
| Chile | 14.7 | 17.2 | 15.7 | 14.7 | 6.3 | 11.2 | 2.6 | 17.7 |  | 9.9 | 13.5 | 14.8 | 13.5 | 8.2 | 12.0 | 4.5 | 23.4 |
| Chinese Taipei | 18.5 | 14.0 | 14.9 | 9.4 | 3.8 | 15.4 | 3.2 | 20.8 |  | 13.1 | 8.4 | 11.7 | 8.1 | 3.9 | 17.3 | 4.4 | 33.1 |
| Colombia | 26.0 | 20.5 | 14.3 | 9.2 | 4.6 | 9.6 | 2.5 | 13.3 |  | 21.6 | 20.9 | 13.8 | 11.1 | 5.6 | 9.6 | 2.8 | 14.7 |
| Costa Rica | 17.8 | 21.2 | 17.4 | 13.6 | 5.9 | 10.0 | 2.6 | 11.5 |  | 11.4 | 19.2 | 17.2 | 13.1 | 8.8 | 10.5 | 3.3 | 16.5 |
| Croatia | 12.7 | 14.3 | 15.3 | 11.9 | 6.6 | 11.3 | 3.9 | 24.0 |  | 11.4 | 12.3 | 13.4 | 10.9 | 7.9 | 11.1 | 5.5 | 27.6 |
| Czech Republic | 5.2 | 10.0 | 11.8 | 13.0 | 9.6 | 11.0 | 5.3 | 34.1 |  | 8.1 | 14.2 | 12.2 | 12.2 | 7.9 | 10.4 | 4.2 | 31.0 |
| Denmark | 5.5 | 8.5 | 10.3 | 11.6 | 8.7 | 16.6 | 7.9 | 30.9 |  | 7.5 | 7.0 | 7.6 | 8.1 | 7.9 | 18.0 | 8.5 | 35.3 |
| Dominican Republic | 16.1 | 17.8 | 17.6 | 11.0 | 7.2 | 12.3 | 1.9 | 16.0 |  | 11.8 | 16.4 | 18.1 | 10.1 | 7.0 | 14.0 | 5.1 | 17.5 |
| Estonia | 9.4 | 9.0 | 15.6 | 15.8 | 11.7 | 12.8 | 5.4 | 20.3 |  | 10.0 | 9.5 | 15.5 | 14.9 | 10.2 | 13.5 | 5.5 | 21.1 |
| Finland | 4.2 | 8.7 | 11.1 | 14.3 | 12.4 | 15.8 | 9.6 | 23.9 |  | 6.7 | 8.6 | 11.0 | 12.6 | 10.6 | 15.7 | 8.6 | 26.1 |
| France | 14.3 | 15.1 | 14.2 | 9.7 | 7.4 | 7.7 | 4.8 | 26.8 |  | 12.0 | 12.9 | 13.0 | 12.0 | 9.5 | 9.7 | 4.1 | 26.8 |
| Germany | 5.0 | 8.8 | 10.9 | 10.8 | 8.7 | 12.8 | 6.0 | 37.0 |  | 6.7 | 8.0 | 9.4 | 10.0 | 7.8 | 14.5 | 6.3 | 37.3 |
| Greece | 15.0 | 14.6 | 17.6 | 15.4 | 9.0 | 8.5 | 3.7 | 16.1 |  | 12.6 | 12.2 | 15.0 | 15.1 | 9.8 | 10.5 | 4.7 | 20.1 |
| Hong Kong | 20.7 | 15.3 | 13.9 | 11.2 | 4.8 | 11.9 | 3.5 | 18.7 |  | 14.6 | 14.7 | 11.8 | 10.2 | 5.5 | 12.2 | 3.6 | 27.4 |
| Hungary | 8.4 | 9.0 | 12.2 | 12.3 | 8.5 | 14.1 | 5.0 | 30.4 |  | 8.8 | 8.4 | 11.4 | 12.8 | 6.6 | 14.8 | 5.2 | 32.1 |
| Iceland | 8.6 | 8.4 | 10.7 | 12.5 | 11.2 | 12.7 | 11.3 | 24.6 |  | 8.7 | 9.3 | 9.9 | 9.3 | 10.1 | 13.5 | 10.1 | 29.0 |
| Ireland | 10.9 | 16.5 | 17.6 | 16.3 | 10.0 | 11.2 | 5.3 | 12.2 |  | 8.5 | 11.0 | 14.4 | 12.2 | 9.7 | 13.8 | 7.9 | 22.5 |
| Israel | 22.6 | 14.2 | 13.8 | 10.9 | 6.1 | 5.0 | 18.4 | 8.9 |  | 15.8 | 13.4 | 13.4 | 13.0 | 9.4 | 8.3 | 10.0 | 16.7 |
| Japan | 30.6 | 7.7 | 5.8 | 5.9 | 3.0 | 16.5 | 10.5 | 20.0 |  | 23.5 | 6.1 | 5.5 | 5.3 | 3.3 | 13.2 | 10.8 | 32.3 |
| Korea | 27.4 | 14.4 | 13.7 | 9.0 | 5.0 | 13.6 | 3.4 | 13.5 |  | 12.9 | 10.3 | 13.5 | 12.1 | 5.1 | 15.5 | 4.5 | 26.1 |
| Latvia | 5.8 | 8.4 | 12.2 | 14.1 | 10.5 | 12.5 | 6.3 | 30.1 |  | 8.7 | 9.7 | 12.9 | 13.5 | 9.2 | 12.8 | 5.6 | 27.5 |
| Lithuania | 7.7 | 10.1 | 12.5 | 14.2 | 8.1 | 12.3 | 5.2 | 29.9 |  | 10.0 | 9.9 | 11.9 | 11.7 | 9.2 | 13.0 | 4.6 | 29.7 |
| Luxembourg | 13.6 | 17.7 | 16.4 | 12.2 | 8.6 | 8.3 | 4.0 | 19.3 |  | 11.5 | 13.9 | 13.5 | 11.9 | 10.3 | 11.1 | 5.3 | 22.6 |
| Mexico | 11.4 | 15.3 | 16.9 | 15.9 | 7.7 | 14.0 | 3.0 | 15.8 |  | 8.5 | 17.3 | 17.4 | 13.3 | 8.3 | 11.9 | 4.7 | 18.5 |
| Montenegro | 6.5 | 9.8 | 14.6 | 16.7 | 11.1 | 13.3 | 4.2 | 23.8 |  | 5.2 | 8.5 | 13.9 | 13.0 | 10.2 | 12.2 | 5.9 | 31.1 |
| Netherlands | 6.0 | 7.2 | 8.4 | 8.0 | 6.0 | 24.6 | 11.1 | 28.7 |  | 6.1 | 6.8 | 8.5 | 8.0 | 4.5 | 25.1 | 12.1 | 28.9 |
| New Zealand | 10.7 | 12.0 | 13.7 | 13.8 | 10.1 | 15.7 | 6.4 | 17.7 |  | 10.5 | 8.7 | 11.0 | 11.5 | 10.2 | 15.6 | 7.7 | 24.9 |
| Norway | 5.6 | 6.8 | 8.8 | 10.9 | 9.0 | 16.9 | 9.0 | 33.0 |  | 8.6 | 7.8 | 9.5 | 8.6 | 6.4 | 16.4 | 6.5 | 36.2 |
| Peru | 9.9 | 21.8 | 16.8 | 15.0 | 7.6 | 10.7 | 2.7 | 15.5 |  | 7.8 | 20.2 | 18.2 | 13.1 | 6.9 | 11.0 | 3.0 | 19.8 |
| Poland | 5.9 | 8.7 | 10.3 | 10.8 | 10.1 | 11.6 | 6.9 | 35.7 |  | 8.1 | 7.3 | 8.4 | 10.5 | 8.5 | 10.7 | 6.8 | 39.6 |
| Portugal | 16.2 | 14.2 | 18.7 | 12.1 | 6.5 | 9.7 | 2.8 | 19.8 |  | 14.3 | 11.5 | 15.5 | 12.0 | 7.2 | 9.9 | 3.4 | 26.2 |
| Qatar | 26.6 | 17.7 | 14.7 | 13.4 | 7.1 | 6.8 | 2.2 | 11.5 |  | 19.4 | 15.8 | 14.7 | 12.8 | 9.2 | 9.1 | 3.6 | 15.5 |
| Russian Federation | 6.1 | 9.4 | 13.4 | 15.7 | 10.2 | 9.2 | 7.4 | 28.6 |  | 6.6 | 7.4 | 12.2 | 17.8 | 10.4 | 8.2 | 7.9 | 29.5 |
| Singapore | 17.3 | 15.9 | 15.0 | 9.3 | 4.7 | 11.8 | 2.8 | 23.2 |  | 13.3 | 14.0 | 13.8 | 10.7 | 5.3 | 11.4 | 3.3 | 28.3 |
| Slovakia | 6.4 | 12.1 | 13.4 | 12.1 | 9.4 | 12.3 | 4.3 | 30.0 |  | 9.8 | 10.0 | 12.7 | 11.4 | 8.3 | 12.0 | 4.0 | 31.7 |
| Slovenia | 5.8 | 13.1 | 16.1 | 16.0 | 9.2 | 11.9 | 5.5 | 22.4 |  | 6.8 | 12.4 | 12.6 | 13.5 | 9.4 | 13.2 | 5.9 | 26.1 |
| Spain | 15.8 | 12.9 | 18.9 | 14.4 | 8.8 | 10.3 | 3.7 | 15.2 |  | 17.4 | 10.5 | 14.6 | 12.4 | 9.0 | 10.6 | 4.1 | 21.4 |
| Sweden | 7.1 | 9.3 | 10.4 | 11.2 | 9.4 | 14.7 | 7.9 | 30.0 |  | 10.4 | 9.3 | 10.2 | 10.6 | 8.3 | 15.4 | 6.4 | 29.4 |
| Switzerland | 7.2 | 12.8 | 11.9 | 9.9 | 6.7 | 12.6 | 6.5 | 32.2 |  | 7.6 | 10.3 | 11.0 | 11.0 | 7.7 | 13.2 | 6.7 | 32.5 |
| Thailand | 4.5 | 14.8 | 17.3 | 16.1 | 6.1 | 15.8 | 1.8 | 23.6 |  | 4.9 | 17.1 | 14.8 | 14.9 | 6.9 | 13.8 | 1.9 | 25.6 |
| United Arab Emirates | 26.1 | 21.8 | 17.7 | 11.3 | 6.0 | 3.6 | 3.5 | 10.0 |  | 14.0 | 20.2 | 17.8 | 16.4 | 8.7 | 6.2 | 4.4 | 12.2 |
| Tunisia | 19.6 | 19.1 | 15.7 | 9.7 | 5.9 | 9.6 | 1.4 | 18.9 |  | 15.8 | 20.3 | 15.3 | 12.0 | 6.4 | 8.2 | 1.8 | 20.2 |
| Turkey | 32.4 | 16.7 | 14.4 | 10.0 | 6.2 | 7.2 | 1.9 | 11.1 |  | 21.7 | 17.0 | 14.5 | 11.8 | 7.6 | 8.2 | 3.1 | 16.0 |
| United Kingdom | 12.5 | 16.3 | 14.0 | 10.8 | 7.4 | 13.3 | 4.3 | 21.5 |  | 10.4 | 12.4 | 11.9 | 11.9 | 8.7 | 13.7 | 5.6 | 25.4 |
| United States | 12.6 | 8.8 | 12.2 | 13.5 | 8.8 | 15.6 | 6.0 | 22.4 |  | 8.8 | 5.7 | 7.9 | 9.3 | 8.1 | 17.4 | 8.0 | 34.7 |
| Uruguay | 18.8 | 12.9 | 17.6 | 12.5 | 7.3 | 11.7 | 4.8 | 14.5 |  | 12.6 | 11.5 | 14.5 | 11.6 | 7.9 | 13.3 | 6.8 | 21.7 |

NB: percent of pupils in each category

| **(d) Out of school vigorous activity (20mins+), days/week** | | | | | | | | | | | | | | | | | |
| --- | --- | --- | --- | --- | --- | --- | --- | --- | --- | --- | --- | --- | --- | --- | --- | --- | --- |
|  | Female | |  |  |  |  |  |  |  | Male |  |  |  |  |  |  |  |
| Country | 0 | 1 | 2 | 3 | 4 | 5 | 6 | 7 |  | 0 | 1 | 2 | 3 | 4 | 5 | 6 | 7 |
| Australia | 20.7 | 16.3 | 19.3 | 15.9 | 10.4 | 8.0 | 3.7 | 5.7 |  | 12.9 | 11.7 | 14.5 | 16.0 | 13.1 | 11.6 | 5.7 | 14.4 |
| Austria | 26.3 | 22.2 | 19.5 | 13.9 | 7.8 | 4.9 | 2.0 | 3.4 |  | 15.4 | 16.1 | 16.3 | 15.8 | 12.6 | 9.4 | 4.8 | 9.7 |
| Belgium | 24.1 | 22.4 | 18.7 | 14.0 | 7.7 | 5.9 | 2.5 | 4.6 |  | 13.8 | 15.7 | 16.3 | 17.5 | 12.5 | 10.4 | 4.8 | 9.1 |
| Brazil | 46.5 | 15.6 | 12.0 | 8.3 | 4.3 | 6.6 | 1.6 | 5.1 |  | 22.7 | 16.8 | 15.1 | 11.6 | 7.2 | 9.0 | 4.4 | 13.0 |
| Bulgaria | 22.1 | 17.8 | 18.9 | 14.6 | 8.1 | 7.0 | 3.6 | 7.9 |  | 13.1 | 14.5 | 15.6 | 14.5 | 9.1 | 11.3 | 5.9 | 16.0 |
| Canada | 18.4 | 14.5 | 16.5 | 15.0 | 11.6 | 11.6 | 4.7 | 7.7 |  | 11.8 | 9.0 | 12.7 | 14.1 | 11.5 | 15.1 | 6.9 | 18.9 |
| Chile | 30.3 | 22.6 | 18.8 | 12.5 | 5.6 | 4.1 | 1.5 | 4.6 |  | 12.7 | 17.1 | 16.7 | 16.0 | 11.7 | 8.9 | 4.5 | 12.4 |
| Chinese Taipei | 24.1 | 19.6 | 27.9 | 10.9 | 4.5 | 6.5 | 1.8 | 4.7 |  | 12.8 | 11.6 | 22.7 | 13.6 | 8.0 | 10.4 | 4.0 | 16.8 |
| Colombia | 23.9 | 26.3 | 17.5 | 11.4 | 5.9 | 7.3 | 2.3 | 5.4 |  | 14.2 | 21.9 | 16.2 | 12.5 | 9.6 | 9.1 | 4.9 | 11.6 |
| Costa Rica | 41.1 | 21.9 | 14.1 | 10.3 | 3.6 | 3.6 | 1.9 | 3.5 |  | 14.1 | 18.2 | 16.7 | 16.4 | 10.2 | 8.9 | 4.0 | 11.6 |
| Croatia | 25.5 | 18.9 | 19.0 | 12.6 | 7.4 | 6.3 | 2.6 | 7.7 |  | 13.1 | 12.6 | 14.7 | 13.9 | 10.4 | 11.7 | 7.1 | 16.6 |
| Czech Republic | 12.4 | 18.1 | 17.1 | 18.0 | 11.8 | 9.0 | 4.8 | 8.8 |  | 10.5 | 13.9 | 15.3 | 16.1 | 13.2 | 10.8 | 6.3 | 13.9 |
| Denmark | 12.1 | 14.1 | 16.5 | 20.0 | 12.7 | 11.1 | 5.0 | 8.6 |  | 10.1 | 8.9 | 13.1 | 15.1 | 13.5 | 14.0 | 7.7 | 17.7 |
| Dominican Republic | 21.8 | 20.4 | 18.4 | 12.6 | 7.7 | 8.7 | 2.7 | 7.7 |  | 11.5 | 15.1 | 15.7 | 12.8 | 9.7 | 12.8 | 6.3 | 16.1 |
| Estonia | 16.0 | 14.2 | 19.1 | 16.7 | 11.7 | 10.3 | 4.2 | 7.8 |  | 10.2 | 12.2 | 17.1 | 17.1 | 13.0 | 12.2 | 5.4 | 12.9 |
| Finland | 11.5 | 16.2 | 19.8 | 17.8 | 11.9 | 11.2 | 7.0 | 4.5 |  | 12.7 | 14.1 | 17.1 | 16.1 | 11.7 | 11.5 | 7.8 | 9.0 |
| France | 28.8 | 24.7 | 19.7 | 11.4 | 6.5 | 4.0 | 1.7 | 3.2 |  | 17.3 | 17.5 | 18.5 | 17.3 | 10.4 | 6.3 | 3.7 | 9.0 |
| Germany | 13.4 | 16.8 | 23.4 | 19.2 | 12.1 | 7.5 | 2.5 | 5.0 |  | 10.1 | 11.4 | 16.7 | 18.9 | 15.2 | 12.0 | 5.4 | 10.4 |
| Greece | 24.9 | 17.6 | 18.2 | 14.0 | 8.1 | 7.4 | 3.8 | 6.1 |  | 13.1 | 11.6 | 14.7 | 16.5 | 12.3 | 11.6 | 5.8 | 14.3 |
| Hong Kong | 27.7 | 26.3 | 18.2 | 12.6 | 4.9 | 4.9 | 1.5 | 3.8 |  | 17.7 | 20.2 | 16.4 | 13.3 | 7.3 | 9.3 | 3.4 | 12.4 |
| Hungary | 17.1 | 13.6 | 19.9 | 17.0 | 10.3 | 9.0 | 3.6 | 9.5 |  | 10.4 | 9.6 | 13.6 | 16.2 | 11.8 | 14.3 | 6.6 | 17.5 |
| Iceland | 11.1 | 9.3 | 13.3 | 11.7 | 13.3 | 15.0 | 11.2 | 15.2 |  | 8.2 | 7.9 | 9.1 | 11.9 | 10.9 | 14.7 | 11.6 | 25.8 |
| Ireland | 20.7 | 16.8 | 18.9 | 16.1 | 9.6 | 7.9 | 4.4 | 5.7 |  | 8.3 | 9.3 | 13.5 | 15.4 | 15.4 | 14.5 | 8.4 | 15.2 |
| Israel | 26.3 | 17.7 | 14.8 | 9.8 | 6.1 | 4.2 | 17.2 | 3.9 |  | 13.4 | 12.1 | 15.9 | 15.4 | 12.1 | 9.4 | 10.5 | 11.2 |
| Japan | 43.1 | 13.1 | 9.8 | 8.0 | 3.0 | 5.6 | 8.3 | 9.2 |  | 24.2 | 9.0 | 7.9 | 8.3 | 4.4 | 7.9 | 13.4 | 24.9 |
| Korea | 41.6 | 18.9 | 18.6 | 9.5 | 2.7 | 4.4 | 1.1 | 3.2 |  | 15.1 | 13.8 | 21.3 | 15.3 | 8.6 | 9.7 | 3.8 | 12.5 |
| Latvia | 15.8 | 14.9 | 19.2 | 16.8 | 11.5 | 10.7 | 4.1 | 7.1 |  | 9.1 | 11.2 | 15.0 | 17.6 | 12.4 | 14.0 | 6.5 | 14.2 |
| Lithuania | 18.4 | 16.1 | 19.7 | 16.8 | 9.9 | 8.7 | 3.8 | 6.6 |  | 10.0 | 9.3 | 15.7 | 16.1 | 12.8 | 14.7 | 6.1 | 15.3 |
| Luxembourg | 21.0 | 19.0 | 18.5 | 15.8 | 9.7 | 6.5 | 3.3 | 6.1 |  | 12.0 | 12.6 | 15.2 | 15.3 | 13.0 | 10.9 | 6.2 | 14.9 |
| Mexico | 18.7 | 20.3 | 20.4 | 13.6 | 8.2 | 9.6 | 2.9 | 6.4 |  | 11.0 | 16.4 | 17.4 | 13.5 | 8.7 | 13.3 | 6.0 | 13.7 |
| Montenegro | 21.0 | 16.0 | 17.0 | 15.1 | 7.9 | 8.5 | 4.5 | 10.1 |  | 8.3 | 11.1 | 13.4 | 13.3 | 10.1 | 12.6 | 6.3 | 25.0 |
| Netherlands | 19.3 | 16.7 | 18.2 | 21.1 | 11.6 | 7.1 | 3.2 | 2.8 |  | 12.9 | 12.6 | 14.0 | 25.1 | 15.0 | 8.9 | 5.4 | 6.1 |
| New Zealand | 21.8 | 13.1 | 19.2 | 14.4 | 11.7 | 8.8 | 5.0 | 6.0 |  | 16.7 | 10.5 | 14.4 | 14.3 | 12.6 | 12.5 | 7.0 | 12.1 |
| Norway | 11.4 | 12.8 | 16.0 | 19.9 | 14.3 | 11.3 | 6.1 | 8.3 |  | 10.5 | 11.0 | 14.0 | 14.1 | 12.4 | 14.3 | 8.0 | 15.7 |
| Peru | 22.1 | 28.0 | 20.4 | 12.3 | 6.1 | 4.9 | 2.1 | 4.1 |  | 7.3 | 20.0 | 19.0 | 16.6 | 10.2 | 9.5 | 4.5 | 12.8 |
| Poland | 14.0 | 14.2 | 14.8 | 16.8 | 10.7 | 9.0 | 6.1 | 14.3 |  | 8.8 | 9.1 | 11.4 | 14.3 | 12.9 | 12.3 | 7.1 | 24.1 |
| Portugal | 27.3 | 17.5 | 21.8 | 12.8 | 7.8 | 5.6 | 2.3 | 5.0 |  | 15.2 | 12.1 | 17.6 | 15.5 | 13.5 | 9.9 | 4.2 | 12.0 |
| Qatar | 33.4 | 19.1 | 16.2 | 11.1 | 6.8 | 5.5 | 2.4 | 5.5 |  | 16.4 | 16.8 | 15.9 | 13.2 | 9.7 | 9.9 | 5.0 | 13.3 |
| Russian Federation | 15.7 | 14.7 | 16.4 | 21.4 | 10.7 | 8.2 | 4.6 | 8.3 |  | 8.8 | 10.5 | 14.1 | 21.4 | 11.7 | 9.2 | 7.4 | 16.9 |
| Singapore | 22.9 | 27.4 | 24.2 | 13.8 | 5.8 | 2.7 | 1.1 | 2.1 |  | 14.8 | 19.8 | 20.1 | 15.9 | 9.7 | 9.3 | 2.9 | 7.6 |
| Slovakia | 14.6 | 18.9 | 19.0 | 17.4 | 9.3 | 7.6 | 4.0 | 9.2 |  | 10.5 | 11.9 | 14.7 | 15.5 | 11.0 | 11.7 | 5.6 | 19.1 |
| Slovenia | 13.6 | 18.1 | 18.7 | 16.6 | 10.3 | 8.1 | 5.1 | 9.5 |  | 9.0 | 11.4 | 13.0 | 13.8 | 12.6 | 13.2 | 8.0 | 19.0 |
| Spain | 26.6 | 15.3 | 22.6 | 13.5 | 9.6 | 6.2 | 2.9 | 3.1 |  | 16.7 | 10.9 | 16.9 | 16.3 | 15.8 | 9.2 | 4.7 | 9.5 |
| Sweden | 12.5 | 14.0 | 16.9 | 17.2 | 15.3 | 10.1 | 6.8 | 7.2 |  | 12.2 | 11.0 | 13.5 | 13.4 | 11.8 | 14.4 | 8.2 | 15.6 |
| Switzerland | 14.9 | 19.7 | 20.5 | 17.5 | 11.2 | 8.6 | 3.0 | 4.7 |  | 9.7 | 11.9 | 16.5 | 19.7 | 14.1 | 11.9 | 6.1 | 10.2 |
| Thailand | 13.5 | 27.4 | 24.2 | 16.5 | 5.7 | 6.4 | 1.2 | 5.2 |  | 9.1 | 19.6 | 16.9 | 14.9 | 7.1 | 13.4 | 2.4 | 16.7 |
| United Arab Emirates | 36.1 | 21.8 | 15.6 | 11.2 | 5.8 | 3.1 | 2.0 | 4.5 |  | 12.9 | 18.9 | 19.2 | 14.9 | 10.6 | 7.3 | 4.4 | 11.8 |
| Tunisia | 30.7 | 22.7 | 18.4 | 11.5 | 4.6 | 4.9 | 1.1 | 6.1 |  | 12.8 | 20.0 | 19.2 | 14.7 | 8.8 | 8.0 | 3.3 | 13.2 |
| Turkey | 34.2 | 21.4 | 15.2 | 10.6 | 5.8 | 5.5 | 1.6 | 5.7 |  | 17.4 | 17.8 | 16.0 | 12.2 | 8.9 | 9.6 | 4.4 | 13.7 |
| United Kingdom | 28.1 | 24.0 | 18.0 | 12.6 | 6.6 | 4.6 | 2.7 | 3.4 |  | 14.9 | 15.0 | 17.3 | 15.9 | 11.3 | 11.2 | 4.5 | 10.0 |
| United States | 22.0 | 12.3 | 14.1 | 13.6 | 8.4 | 14.2 | 6.4 | 8.9 |  | 11.4 | 7.3 | 9.6 | 10.9 | 9.0 | 18.7 | 8.7 | 24.3 |
| Uruguay | 33.2 | 14.0 | 19.0 | 14.2 | 5.8 | 6.8 | 3.1 | 3.9 |  | 14.0 | 10.4 | 15.4 | 15.4 | 10.3 | 11.9 | 8.9 | 13.7 |

NB: percent of pupils in each category

Supplementary Table 5. Tabulations of physical activity outcomes by wealth (top and bottom quintile).

| **(a) Physical activity in school, classes per week** | | | | | | | | | | | | | | | | | |
| --- | --- | --- | --- | --- | --- | --- | --- | --- | --- | --- | --- | --- | --- | --- | --- | --- | --- |
|  | Wealth Q1 | | | | | | |  |  | Wealth Q5 | |  |  |  |  |  |  |
| Country | 0 | 1 | 2 | 3 | 4 | 5 | 6 | 7 |  | 0 | 1 | 2 | 3 | 4 | 5 | 6 | 7 |
| Australia | 16.8 | 15.7 | 25.2 | 21.1 | 11.2 | 9.9 | 0.0 | 0.0 |  | 13.6 | 17.9 | 28.4 | 21.9 | 9.2 | 9.0 | 0.0 | 0.0 |
| Austria | 10.4 | 63.3 | 17.2 | 4.1 | 1.3 | 2.2 | 1.5 | 0.0 |  | 11.5 | 64.6 | 14.4 | 3.4 | 1.7 | 2.0 | 2.4 | 0.0 |
| Belgium | 1.9 | 53.2 | 37.4 | 3.3 | 1.9 | 2.0 | 0.0 | 0.4 |  | 1.0 | 48.5 | 41.8 | 3.0 | 2.0 | 3.0 | 0.2 | 0.6 |
| Brazil | 11.2 | 37.5 | 33.9 | 3.3 | 2.1 | 3.7 | 1.6 | 6.6 |  | 13.6 | 41.5 | 31.1 | 3.0 | 1.5 | 2.3 | 1.1 | 5.7 |
| Bulgaria | 3.1 | 4.5 | 48.8 | 27.5 | 3.7 | 4.2 | 1.5 | 6.8 |  | 2.2 | 6.2 | 48.8 | 25.0 | 4.5 | 3.8 | 1.9 | 7.6 |
| Canada | 22.9 | 18.2 | 10.2 | 12.9 | 4.9 | 31.0 | 0.0 | 0.0 |  | 23.2 | 5.8 | 8.1 | 15.2 | 5.2 | 42.6 | 0.0 | 0.0 |
| Chile | 1.3 | 53.8 | 28.7 | 2.3 | 1.7 | 12.3 | 0.0 | 0.0 |  | 1.0 | 49.8 | 37.9 | 2.0 | 2.0 | 7.3 | 0.0 | 0.0 |
| Chinese Taipei | 1.8 | 22.0 | 69.6 | 3.5 | 0.2 | 2.9 | 0.0 | 0.0 |  | 0.7 | 24.3 | 70.0 | 2.4 | 0.7 | 1.9 | 0.0 | 0.0 |
| Colombia | 3.1 | 59.3 | 20.6 | 1.5 | 1.3 | 7.3 | 6.9 | 0.0 |  | 2.8 | 67.5 | 20.9 | 1.3 | 1.3 | 2.0 | 4.1 | 0.0 |
| Costa Rica | 14.0 | 81.4 | 2.9 | 0.1 | 0.1 | 1.5 | 0.0 | 0.0 |  | 3.8 | 93.8 | 1.6 | 0.0 | 0.0 | 0.8 | 0.0 | 0.0 |
| Croatia | 1.0 | 37.9 | 58.8 | 1.0 | 0.5 | 0.4 | 0.5 | 0.0 |  | 1.2 | 32.2 | 64.6 | 0.9 | 0.7 | 0.2 | 0.2 | 0.0 |
| Czech Republic | 3.1 | 37.9 | 47.9 | 5.7 | 2.3 | 3.1 | 0.0 | 0.0 |  | 3.5 | 41.9 | 44.6 | 4.5 | 1.7 | 3.7 | 0.0 | 0.0 |
| Denmark | 3.6 | 72.2 | 13.9 | 3.8 | 2.9 | 3.6 | 0.0 | 0.0 |  | 2.1 | 70.7 | 13.8 | 4.4 | 2.9 | 6.1 | 0.0 | 0.0 |
| Dominican Republic | 9.6 | 23.0 | 31.9 | 8.3 | 1.8 | 25.4 | 0.0 | 0.0 |  | 8.0 | 26.6 | 36.0 | 5.5 | 3.4 | 20.5 | 0.0 | 0.0 |
| Estonia | 2.4 | 31.7 | 61.1 | 1.7 | 0.6 | 2.6 | 0.0 | 0.0 |  | 4.3 | 31.2 | 59.8 | 1.6 | 0.5 | 2.6 | 0.0 | 0.0 |
| Finland | 1.3 | 50.1 | 33.6 | 6.8 | 2.3 | 5.9 | 0.0 | 0.0 |  | 1.4 | 44.3 | 35.1 | 9.3 | 3.9 | 6.1 | 0.0 | 0.0 |
| France | 3.9 | 65.9 | 20.0 | 4.7 | 1.0 | 4.5 | 0.0 | 0.0 |  | 3.9 | 70.4 | 16.5 | 4.2 | 0.9 | 4.2 | 0.0 | 0.0 |
| Germany | 1.4 | 64.9 | 26.5 | 3.2 | 1.5 | 1.6 | 0.9 | 0.0 |  | 2.3 | 68.5 | 23.1 | 3.0 | 0.9 | 1.3 | 0.9 | 0.0 |
| Greece | 3.0 | 6.6 | 70.4 | 5.8 | 2.3 | 2.6 | 0.8 | 8.4 |  | 3.4 | 6.4 | 72.1 | 5.9 | 1.5 | 2.3 | 1.0 | 7.5 |
| Hong Kong | 1.7 | 90.6 | 5.9 | 0.5 | 0.2 | 1.2 | 0.0 | 0.0 |  | 1.2 | 87.3 | 9.0 | 0.6 | 0.5 | 1.4 | 0.0 | 0.0 |
| Hungary | 1.7 | 2.3 | 12.4 | 32.0 | 18.5 | 31.3 | 0.4 | 1.3 |  | 0.7 | 1.2 | 8.7 | 34.4 | 19.6 | 33.4 | 0.7 | 1.4 |
| Iceland | 2.6 | 14.4 | 51.8 | 14.2 | 8.2 | 2.7 | 1.7 | 4.5 |  | 2.7 | 10.9 | 54.3 | 16.6 | 5.8 | 2.6 | 1.4 | 5.7 |
| Ireland | 9.4 | 75.9 | 10.3 | 1.2 | 0.5 | 2.7 | 0.0 | 0.0 |  | 9.8 | 74.1 | 11.6 | 0.8 | 0.8 | 2.9 | 0.0 | 0.0 |
| Israel | 11.0 | 27.8 | 40.9 | 4.4 | 3.2 | 3.2 | 1.4 | 8.0 |  | 6.9 | 21.6 | 53.6 | 3.0 | 2.0 | 2.8 | 1.5 | 8.6 |
| Japan | 0.1 | 5.3 | 39.6 | 51.2 | 2.7 | 0.5 | 0.7 | 0.0 |  | 0.1 | 4.7 | 41.9 | 50.4 | 2.5 | 0.3 | 0.1 | 0.0 |
| Korea | 0.7 | 7.9 | 73.4 | 12.9 | 2.7 | 2.5 | 0.0 | 0.0 |  | 0.3 | 6.2 | 70.3 | 16.0 | 4.8 | 2.4 | 0.0 | 0.0 |
| Latvia | 5.0 | 13.3 | 66.3 | 8.0 | 1.1 | 6.3 | 0.0 | 0.0 |  | 6.5 | 15.3 | 61.1 | 6.3 | 2.1 | 8.6 | 0.0 | 0.0 |
| Lithuania | 7.6 | 6.8 | 75.0 | 4.2 | 2.0 | 4.4 | 0.0 | 0.0 |  | 7.1 | 5.4 | 71.4 | 5.1 | 2.0 | 9.0 | 0.0 | 0.0 |
| Luxembourg | 3.8 | 56.4 | 29.8 | 5.5 | 1.4 | 3.1 | 0.0 | 0.0 |  | 2.3 | 56.6 | 26.9 | 5.1 | 3.6 | 5.5 | 0.0 | 0.0 |
| Mexico | 21.9 | 30.3 | 30.9 | 7.0 | 1.6 | 3.1 | 5.3 | 0.0 |  | 21.6 | 28.1 | 38.5 | 4.2 | 1.8 | 1.6 | 4.3 | 0.0 |
| Montenegro | 2.5 | 5.6 | 56.5 | 8.2 | 2.4 | 3.9 | 2.1 | 18.7 |  | 4.7 | 4.8 | 49.7 | 10.4 | 3.9 | 5.4 | 2.2 | 19.0 |
| Netherlands | 3.9 | 58.6 | 29.6 | 5.3 | 1.7 | 0.9 | 0.0 | 0.0 |  | 4.3 | 61.7 | 26.5 | 4.6 | 1.6 | 1.3 | 0.0 | 0.0 |
| New Zealand | 44.3 | 5.1 | 10.2 | 9.3 | 16.3 | 12.1 | 2.7 | 0.0 |  | 35.5 | 8.7 | 12.1 | 9.9 | 19.1 | 10.2 | 4.4 | 0.0 |
| Norway | 0.5 | 40.3 | 46.3 | 9.4 | 1.8 | 0.7 | 0.3 | 0.7 |  | 0.7 | 31.5 | 53.7 | 11.2 | 0.8 | 1.6 | 0.0 | 0.5 |
| Peru | 1.5 | 59.7 | 14.2 | 3.1 | 1.6 | 19.8 | 0.0 | 0.0 |  | 3.9 | 70.9 | 14.8 | 1.4 | 1.0 | 8.0 | 0.0 | 0.0 |
| Poland | 0.1 | 1.0 | 21.7 | 30.3 | 42.3 | 4.6 | 0.0 | 0.0 |  | 1.0 | 1.6 | 22.9 | 32.5 | 34.9 | 7.2 | 0.0 | 0.0 |
| Portugal | 0.7 | 12.7 | 76.7 | 5.2 | 0.8 | 0.6 | 0.5 | 2.9 |  | 0.4 | 6.6 | 82.9 | 6.7 | 1.1 | 0.4 | 0.4 | 1.5 |
| Qatar | 10.0 | 49.7 | 16.6 | 6.1 | 3.6 | 13.9 | 0.0 | 0.0 |  | 17.2 | 33.7 | 16.7 | 10.6 | 5.2 | 16.6 | 0.0 | 0.0 |
| Russian Federation | 2.5 | 3.2 | 15.3 | 66.5 | 4.0 | 1.7 | 1.5 | 5.3 |  | 4.6 | 4.6 | 17.9 | 55.4 | 5.7 | 2.1 | 1.5 | 8.2 |
| Singapore | 2.2 | 32.7 | 55.5 | 5.5 | 1.8 | 2.3 | 0.0 | 0.0 |  | 4.3 | 40.5 | 42.4 | 10.5 | 1.0 | 1.3 | 0.0 | 0.0 |
| Slovakia | 4.3 | 16.5 | 62.7 | 11.1 | 2.4 | 3.0 | 0.0 | 0.0 |  | 4.4 | 14.3 | 63.9 | 10.2 | 2.3 | 5.0 | 0.0 | 0.0 |
| Slovenia | 1.1 | 20.9 | 55.1 | 20.3 | 1.0 | 1.7 | 0.0 | 0.0 |  | 1.1 | 18.2 | 56.2 | 18.7 | 1.6 | 4.2 | 0.0 | 0.0 |
| Spain | 1.7 | 8.6 | 86.4 | 1.6 | 0.3 | 1.4 | 0.0 | 0.0 |  | 1.9 | 9.5 | 86.9 | 0.6 | 0.3 | 0.8 | 0.0 | 0.0 |
| Sweden | 3.0 | 17.9 | 62.5 | 5.4 | 1.6 | 2.1 | 0.8 | 6.8 |  | 1.7 | 15.5 | 60.6 | 6.2 | 3.3 | 1.7 | 0.8 | 10.2 |
| Switzerland | 4.3 | 25.4 | 54.6 | 10.8 | 1.7 | 1.2 | 2.0 | 0.0 |  | 4.4 | 19.7 | 58.1 | 9.5 | 2.0 | 2.6 | 3.6 | 0.0 |
| Thailand | 9.3 | 70.1 | 10.8 | 2.3 | 1.1 | 6.3 | 0.0 | 0.0 |  | 5.4 | 81.0 | 8.1 | 1.2 | 0.8 | 3.5 | 0.0 | 0.0 |
| United Arab Emirates | 5.8 | 30.6 | 39.0 | 10.8 | 3.1 | 10.6 | 0.0 | 0.0 |  | 12.6 | 42.2 | 26.6 | 6.6 | 3.3 | 8.7 | 0.0 | 0.0 |
| Tunisia | 4.0 | 46.8 | 21.7 | 2.6 | 3.1 | 21.9 | 0.0 | 0.0 |  | 5.1 | 46.7 | 25.8 | 2.4 | 4.9 | 15.1 | 0.0 | 0.0 |
| Turkey | 8.5 | 47.5 | 24.5 | 4.3 | 2.2 | 13.0 | 0.0 | 0.0 |  | 8.2 | 32.0 | 38.3 | 5.9 | 3.2 | 12.4 | 0.0 | 0.0 |
| United Kingdom | 5.6 | 41.6 | 35.0 | 8.2 | 4.3 | 5.2 | 0.0 | 0.0 |  | 3.7 | 33.4 | 36.5 | 13.3 | 7.8 | 5.3 | 0.0 | 0.0 |
| United States | 40.5 | 3.4 | 6.9 | 11.5 | 4.6 | 33.1 | 0.0 | 0.0 |  | 42.1 | 1.8 | 4.5 | 11.3 | 6.0 | 34.3 | 0.0 | 0.0 |
| Uruguay | 23.0 | 8.8 | 50.7 | 5.7 | 1.3 | 3.8 | 6.8 | 0.0 |  | 14.2 | 17.1 | 53.1 | 5.4 | 1.8 | 3.8 | 4.6 | 0.0 |

NB: percent of pupils in each category

| **(b) Physical activity classes in school, hours per week** | | | | | | | | | | | | | | | | | | | |
| --- | --- | --- | --- | --- | --- | --- | --- | --- | --- | --- | --- | --- | --- | --- | --- | --- | --- | --- | --- |
|  | Wealth Q1 | | |  |  |  | |  | | Wealth Q5 | |  | |  |  | |  | |  |
| Country | 0 hours | | 0-1 hours | 1-2 hours | 2-3 hours | 3-4 hours | | > 4 hours | | 0 hours | | 0-1 hours | | 1-2 hours | 2-3 hours | | 3-4 hours | | > 4 hours |
| Australia | 16.7 | | 12.1 | 25.1 | 20.8 | 14.8 | 10.4 | | 13.6 | | | 14.9 | | 29.3 | 21.2 | | 12.4 | | 8.6 |
| Austria | 10.1 | | 62.1 | 18.6 | 4.5 | 1.3 | 3.4 | | 11.5 | | | 63.6 | | 15.3 | 3.4 | | 1.8 | | 4.3 |
| Belgium | 1.9 | | 53.1 | 37.7 | 3.8 | 1.6 | 1.8 | | 1.0 | | | 48.1 | | 42.1 | 3.4 | | 2.1 | | 3.3 |
| Brazil | 11.4 | | 37.5 | 35.0 | 4.4 | 3.0 | 8.6 | | 13.7 | | | 40.8 | | 32.4 | 3.1 | | 2.3 | | 7.8 |
| Bulgaria | 3.0 | | 5.8 | 63.2 | 15.4 | 5.7 | 7.0 | | 2.1 | | | 6.0 | | 65.5 | 12.3 | | 5.7 | | 8.4 |
| Canada | 22.7 | | 3.4 | 19.1 | 9.7 | 10.9 | 34.3 | | 23.4 | | | 1.5 | | 7.5 | 8.7 | | 12.4 | | 46.6 |
| Chile | 1.3 | | 37.2 | 36.4 | 9.5 | 7.6 | 8.0 | | 1.0 | | | 33.2 | | 43.5 | 12.6 | | 5.1 | | 4.6 |
| Chinese Taipei | 1.8 | | 23.1 | 68.4 | 3.8 | 1.5 | 1.4 | | 0.7 | | | 25.0 | | 69.4 | 2.7 | | 0.8 | | 1.4 |
| Colombia | 2.9 | | 53.2 | 26.9 | 2.6 | 3.4 | 11.0 | | 2.3 | | | 54.2 | | 33.3 | 2.3 | | 1.9 | | 6.1 |
| Costa Rica | 14.1 | | 81.6 | 2.8 | 0.1 | 1.3 | 0.0 | | 3.6 | | | 94.0 | | 1.7 | 0.2 | | 0.4 | | 0.0 |
| Croatia | 1.0 | | 37.0 | 59.6 | 1.6 | 0.4 | 0.4 | | 1.2 | | | 32.0 | | 64.4 | 1.8 | | 0.5 | | 0.2 |
| Czech Republic | 3.0 | | 37.9 | 47.6 | 8.5 | 2.7 | 0.4 | | 3.6 | | | 42.5 | | 44.5 | 5.6 | | 3.1 | | 0.8 |
| Denmark | 3.6 | | 66.4 | 19.6 | 5.2 | 2.9 | 2.3 | | 2.0 | | | 64.2 | | 18.9 | 7.2 | | 4.3 | | 3.4 |
| Dominican Republic | 9.8 | | 23.8 | 30.4 | 10.1 | 9.1 | 16.6 | | 7.8 | | | 26.9 | | 35.0 | 9.3 | | 9.6 | | 11.3 |
| Estonia | 2.4 | | 31.4 | 61.0 | 2.7 | 2.1 | 0.4 | | 4.2 | | | 31.1 | | 59.6 | 2.6 | | 2.1 | | 0.4 |
| Finland | 1.3 | | 46.9 | 33.0 | 12.0 | 4.9 | 1.8 | | 1.2 | | | 42.0 | | 32.3 | 15.5 | | 7.4 | | 1.5 |
| France | 3.6 | | 63.3 | 22.3 | 5.7 | 1.7 | 3.3 | | 3.8 | | | 68.0 | | 18.8 | 4.4 | | 1.4 | | 3.6 |
| Germany | 1.4 | | 60.7 | 27.3 | 7.7 | 1.7 | 1.2 | | 2.3 | | | 64.8 | | 24.3 | 6.2 | | 1.3 | | 1.1 |
| Greece | 2.8 | | 9.7 | 69.7 | 6.2 | 2.8 | 8.9 | | 3.2 | | | 8.4 | | 71.9 | 5.3 | | 2.9 | | 8.3 |
| Hong Kong | 1.7 | | 88.6 | 8.1 | 0.9 | 0.4 | 0.3 | | 1.2 | | | 84.2 | | 12.2 | 1.3 | | 0.8 | | 0.4 |
| Hungary | 1.7 | | 3.4 | 12.8 | 49.4 | 29.9 | 2.8 | | 0.7 | | | 1.3 | | 9.7 | 52.2 | | 33.1 | | 3.0 |
| Iceland | 2.6 | | 14.5 | 61.5 | 10.7 | 5.4 | 5.3 | | 2.8 | | | 10.6 | | 64.0 | 10.1 | | 5.4 | | 7.2 |
| Ireland | 9.4 | | 75.9 | 11.1 | 0.8 | 2.2 | 0.6 | | 9.7 | | | 74.0 | | 12.5 | 1.0 | | 2.6 | | 0.2 |
| Israel | 11.3 | | 27.8 | 41.4 | 6.4 | 3.4 | 9.8 | | 6.9 | | | 21.2 | | 54.4 | 4.5 | | 2.8 | | 10.1 |
| Japan | 0.1 | | 4.2 | 40.1 | 50.3 | 4.3 | 1.1 | | 0.1 | | | 3.6 | | 41.9 | 50.0 | | 4.0 | | 0.4 |
| Korea | 0.7 | | 8.2 | 73.1 | 14.5 | 2.7 | 0.8 | | 0.3 | | | 6.6 | | 70.0 | 19.1 | | 2.3 | | 1.7 |
| Latvia | 4.9 | | 13.8 | 72.8 | 2.3 | 5.9 | 0.3 | | 6.6 | | | 15.0 | | 67.4 | 2.6 | | 7.5 | | 0.9 |
| Lithuania | 7.5 | | 6.6 | 75.4 | 6.1 | 4.3 | 0.0 | | 7.1 | | | 5.3 | | 71.6 | 7.1 | | 8.8 | | 0.0 |
| Luxembourg | 3.7 | | 55.1 | 30.6 | 5.2 | 2.3 | 3.1 | | 2.2 | | | 54.9 | | 28.2 | 5.7 | | 3.9 | | 5.2 |
| Mexico | 22.0 | | 32.3 | 28.6 | 7.6 | 3.0 | 6.5 | | 21.7 | | | 27.2 | | 39.0 | 4.2 | | 2.7 | | 5.3 |
| Montenegro | 2.3 | | 6.0 | 55.9 | 10.3 | 4.9 | 20.6 | | 4.7 | | | 5.0 | | 50.9 | 13.2 | | 5.4 | | 20.8 |
| Netherlands | 3.9 | | 54.6 | 30.3 | 8.4 | 1.8 | 1.0 | | 4.3 | | | 59.1 | | 26.2 | 7.4 | | 2.0 | | 1.1 |
| New Zealand | 44.4 | | 4.3 | 10.4 | 9.3 | 17.4 | 14.3 | | 35.5 | | | 8.0 | | 12.3 | 10.0 | | 18.9 | | 15.2 |
| Norway | 0.4 | | 38.2 | 44.9 | 12.6 | 2.3 | 1.5 | | 0.6 | | | 28.9 | | 51.4 | 15.3 | | 2.1 | | 1.6 |
| Peru | 1.6 | | 57.5 | 17.9 | 4.8 | 9.2 | 9.0 | | 3.9 | | | 66.2 | | 18.9 | 2.1 | | 5.1 | | 3.9 |
| Poland | 0.1 | | 1.1 | 22.3 | 70.9 | 5.2 | 0.3 | | 1.0 | | | 1.7 | | 23.4 | 66.1 | | 7.4 | | 0.5 |
| Portugal | 0.7 | | 8.2 | 43.6 | 38.5 | 4.5 | 4.5 | | 0.4 | | | 4.3 | | 43.6 | 43.2 | | 4.8 | | 3.7 |
| Qatar | 9.6 | | 50.2 | 18.0 | 6.8 | 7.9 | 7.4 | | 17.4 | | | 32.8 | | 17.7 | 10.6 | | 8.7 | | 12.9 |
| Russian Federation | 2.5 | | 3.3 | 34.1 | 51.9 | 1.9 | 6.4 | | 4.6 | | | 4.2 | | 31.2 | 47.7 | | 3.1 | | 9.3 |
| Singapore | 2.2 | | 43.3 | 42.3 | 7.8 | 2.8 | 1.6 | | 4.3 | | | 48.7 | | 35.7 | 6.5 | | 2.3 | | 2.6 |
| Slovakia | 4.1 | | 17.8 | 61.7 | 13.2 | 2.2 | 1.0 | | 4.5 | | | 14.9 | | 63.4 | 11.9 | | 4.3 | | 1.1 |
| Slovenia | 1.1 | | 20.7 | 55.9 | 20.6 | 1.6 | 0.1 | | 1.1 | | | 17.7 | | 57.2 | 19.4 | | 4.1 | | 0.5 |
| Spain | 1.5 | | 10.6 | 83.7 | 2.2 | 0.6 | 1.4 | | 2.0 | | | 9.3 | | 86.1 | 1.4 | | 0.4 | | 0.9 |
| Sweden | 3.0 | | 13.9 | 55.1 | 16.2 | 2.8 | 9.1 | | 1.7 | | | 11.4 | | 49.7 | 18.9 | | 3.8 | | 14.6 |
| Switzerland | 4.3 | | 25.6 | 54.7 | 12.0 | 1.2 | 2.2 | | 4.4 | | | 19.5 | | 58.3 | 11.5 | | 2.6 | | 3.7 |
| Thailand | 9.6 | | 64.2 | 15.6 | 3.2 | 2.2 | 5.2 | | 5.5 | | | 77.9 | | 11.3 | 1.2 | | 1.3 | | 2.9 |
| Tunisia | 6.0 | | 31.0 | 36.5 | 12.2 | 4.6 | 9.7 | | 12.2 | | | 35.2 | | 30.2 | 8.5 | | 4.8 | | 9.1 |
| Turkey | 4.0 | | 48.3 | 22.8 | 4.5 | 19.3 | 1.2 | | 5.1 | | | 46.8 | | 27.6 | 6.6 | | 12.8 | | 1.2 |
| United Arab Emirates | 8.5 | | 47.1 | 25.2 | 5.7 | 7.8 | 5.7 | | 8.2 | | | 31.4 | | 36.7 | 8.3 | | 8.5 | | 6.9 |
| United Kingdom | 5.5 | | 39.5 | 37.3 | 8.7 | 4.6 | 4.4 | | 3.7 | | | 30.5 | | 38.3 | 15.0 | | 7.7 | | 4.9 |
| United States | 40.5 | | 2.7 | 7.3 | 9.3 | 15.4 | 24.9 | | 42.1 | | | 1.1 | | 6.6 | 7.2 | | 17.5 | | 25.4 |
| Uruguay | 23.0 | | 10.7 | 49.6 | 6.8 | 4.2 | 5.7 | | 14.0 | | | 17.2 | | 54.5 | 4.9 | | 6.3 | | 3.1 |

NB: percent of pupils in each category

| **(c) Out of school moderate activity (60mins+), days/week** | | | | | | | | | | | | | | | | | |
| --- | --- | --- | --- | --- | --- | --- | --- | --- | --- | --- | --- | --- | --- | --- | --- | --- | --- |
|  | Wealth Q1 |  |  |  |  |  |  |  |  | Wealth Q5 | |  |  |  |  |  |  |
| Country | 0 | 1 | 2 | 3 | 4 | 5 | 6 | 7 |  | 0 | 1 | 2 | 3 | 4 | 5 | 6 | 7 |
| Australia | 14.7 | 13.8 | 13.7 | 13.7 | 10.3 | 14.7 | 4.3 | 14.8 |  | 8.4 | 9.6 | 13.2 | 14.9 | 9.8 | 13.6 | 6.7 | 23.9 |
| Austria | 11.4 | 14.1 | 10.6 | 8.8 | 5.7 | 13.1 | 4.7 | 31.6 |  | 10.3 | 9.4 | 10.1 | 8.6 | 6.1 | 11.3 | 4.5 | 39.7 |
| Belgium | 17.5 | 17.3 | 12.1 | 9.7 | 6.6 | 11.6 | 3.6 | 21.5 |  | 9.7 | 14.7 | 11.8 | 11.0 | 7.5 | 14.2 | 5.2 | 25.9 |
| Brazil | 22.8 | 22.3 | 17.4 | 11.0 | 5.6 | 7.1 | 2.2 | 11.5 |  | 19.4 | 14.5 | 15.2 | 11.6 | 7.5 | 11.4 | 3.3 | 17.1 |
| Bulgaria | 15.4 | 15.9 | 18.6 | 14.2 | 8.0 | 8.4 | 3.6 | 15.9 |  | 7.9 | 9.8 | 14.2 | 16.7 | 9.4 | 11.7 | 5.4 | 24.9 |
| Canada | 9.8 | 8.8 | 12.0 | 13.8 | 10.2 | 16.3 | 5.2 | 23.9 |  | 4.1 | 4.3 | 9.8 | 13.0 | 10.6 | 18.8 | 6.5 | 33.1 |
| Chile | 12.7 | 18.3 | 15.3 | 14.6 | 7.5 | 10.7 | 3.0 | 18.0 |  | 11.8 | 11.2 | 14.8 | 13.9 | 8.3 | 13.0 | 3.6 | 23.4 |
| Chinese Taipei | 17.0 | 12.8 | 11.8 | 9.1 | 3.0 | 16.7 | 2.8 | 26.9 |  | 13.9 | 9.4 | 13.4 | 9.3 | 3.2 | 17.0 | 4.0 | 29.9 |
| Colombia | 29.0 | 22.4 | 14.4 | 10.0 | 3.9 | 9.1 | 1.4 | 9.9 |  | 20.8 | 17.8 | 12.3 | 10.3 | 5.7 | 9.7 | 3.5 | 20.0 |
| Costa Rica | 16.5 | 22.4 | 18.2 | 12.7 | 7.1 | 9.1 | 2.2 | 11.8 |  | 11.6 | 17.1 | 15.6 | 15.5 | 9.1 | 10.4 | 4.1 | 16.6 |
| Croatia | 14.8 | 14.4 | 16.6 | 13.1 | 5.8 | 10.8 | 4.2 | 20.3 |  | 10.9 | 13.7 | 11.3 | 11.0 | 6.7 | 11.0 | 5.8 | 29.6 |
| Czech Republic | 7.5 | 13.7 | 11.8 | 10.5 | 9.7 | 11.3 | 4.4 | 30.9 |  | 7.0 | 10.0 | 10.7 | 11.3 | 9.6 | 11.0 | 4.2 | 36.1 |
| Denmark | 9.1 | 9.8 | 9.9 | 9.5 | 9.1 | 15.6 | 7.0 | 30.1 |  | 5.7 | 6.5 | 8.3 | 6.9 | 9.5 | 17.8 | 9.2 | 36.2 |
| Dominican Republic | 14.6 | 18.4 | 20.7 | 11.0 | 5.7 | 11.2 | 3.5 | 14.9 |  | 11.4 | 13.0 | 15.9 | 13.1 | 7.6 | 15.4 | 3.6 | 20.0 |
| Estonia | 13.2 | 10.0 | 14.8 | 15.0 | 9.9 | 12.0 | 5.0 | 20.1 |  | 7.7 | 8.4 | 15.5 | 15.7 | 11.3 | 14.0 | 5.8 | 21.5 |
| Finland | 6.7 | 9.8 | 10.8 | 13.2 | 11.7 | 15.7 | 7.9 | 24.3 |  | 4.3 | 6.6 | 12.2 | 14.8 | 10.0 | 14.2 | 8.7 | 29.3 |
| France | 16.6 | 14.1 | 13.0 | 10.8 | 7.9 | 9.0 | 4.9 | 23.6 |  | 12.1 | 13.0 | 12.7 | 9.6 | 9.4 | 9.2 | 3.7 | 30.3 |
| Germany | 7.0 | 10.3 | 11.7 | 10.7 | 9.0 | 13.2 | 5.2 | 33.0 |  | 4.3 | 7.3 | 8.5 | 9.4 | 8.4 | 13.9 | 6.8 | 41.4 |
| Greece | 14.8 | 14.7 | 16.7 | 14.5 | 9.3 | 8.9 | 3.2 | 18.0 |  | 12.1 | 12.2 | 15.2 | 15.3 | 10.7 | 9.8 | 4.0 | 20.9 |
| Hong Kong | 20.9 | 16.4 | 13.2 | 8.8 | 4.5 | 11.7 | 3.5 | 20.9 |  | 14.9 | 13.8 | 13.5 | 10.2 | 6.1 | 11.0 | 3.3 | 27.2 |
| Hungary | 11.5 | 11.8 | 12.1 | 11.4 | 6.5 | 15.2 | 4.6 | 26.9 |  | 6.4 | 5.4 | 10.1 | 11.8 | 8.1 | 14.7 | 5.4 | 38.2 |
| Iceland | 9.8 | 10.9 | 12.1 | 9.4 | 10.3 | 12.0 | 9.3 | 26.3 |  | 6.6 | 8.1 | 8.4 | 10.4 | 10.4 | 12.4 | 12.1 | 31.7 |
| Ireland | 14.0 | 13.3 | 14.8 | 15.0 | 10.5 | 10.9 | 6.1 | 15.6 |  | 7.8 | 11.8 | 14.6 | 15.1 | 9.6 | 13.1 | 8.3 | 19.7 |
| Israel | 26.9 | 17.3 | 14.6 | 11.1 | 7.4 | 6.6 | 3.9 | 12.2 |  | 16.5 | 13.3 | 14.6 | 13.8 | 10.4 | 8.2 | 4.8 | 18.4 |
| Japan | 29.6 | 8.3 | 6.9 | 6.8 | 2.9 | 15.2 | 8.5 | 21.8 |  | 25.1 | 6.9 | 5.0 | 5.0 | 3.5 | 13.7 | 11.0 | 29.7 |
| Korea | 20.8 | 12.7 | 12.1 | 11.6 | 5.0 | 15.6 | 4.4 | 17.8 |  | 18.8 | 12.2 | 13.3 | 10.4 | 4.7 | 13.5 | 2.8 | 24.4 |
| Latvia | 8.6 | 11.9 | 12.5 | 14.7 | 8.2 | 12.1 | 4.1 | 28.0 |  | 5.8 | 6.7 | 10.2 | 12.1 | 11.7 | 13.9 | 5.7 | 33.9 |
| Lithuania | 13.0 | 12.5 | 15.4 | 12.0 | 8.3 | 9.4 | 4.6 | 24.8 |  | 5.4 | 8.0 | 11.3 | 15.5 | 8.5 | 12.1 | 5.0 | 34.3 |
| Luxembourg | 16.7 | 18.4 | 15.0 | 13.0 | 7.5 | 7.1 | 3.6 | 18.7 |  | 9.9 | 12.6 | 12.5 | 12.0 | 12.4 | 11.9 | 5.5 | 23.3 |
| Mexico | 9.9 | 19.5 | 20.3 | 13.9 | 7.2 | 10.1 | 3.0 | 16.1 |  | 9.8 | 12.8 | 15.4 | 15.0 | 8.3 | 14.4 | 5.1 | 19.4 |
| Montenegro | 6.4 | 11.6 | 17.3 | 14.5 | 8.4 | 12.5 | 4.1 | 25.2 |  | 5.8 | 6.6 | 14.4 | 13.2 | 10.4 | 12.6 | 6.3 | 30.9 |
| Netherlands | 9.2 | 8.7 | 10.9 | 6.9 | 5.2 | 24.7 | 11.0 | 23.4 |  | 4.3 | 6.4 | 7.9 | 9.2 | 5.9 | 23.7 | 10.1 | 32.6 |
| New Zealand | 13.1 | 13.4 | 11.6 | 11.9 | 9.2 | 15.3 | 6.0 | 19.5 |  | 8.1 | 7.6 | 11.1 | 11.3 | 11.5 | 14.8 | 8.8 | 26.7 |
| Norway | 9.2 | 10.1 | 11.1 | 10.2 | 7.4 | 15.8 | 7.6 | 28.7 |  | 6.4 | 5.4 | 9.0 | 8.5 | 7.2 | 15.8 | 6.0 | 41.7 |
| Peru | 10.9 | 31.4 | 16.8 | 12.8 | 7.1 | 7.7 | 1.5 | 11.8 |  | 8.7 | 15.1 | 14.7 | 15.3 | 7.7 | 11.9 | 3.9 | 22.9 |
| Poland | 8.8 | 10.1 | 9.3 | 12.8 | 6.7 | 12.3 | 6.4 | 33.6 |  | 6.8 | 6.4 | 8.1 | 9.4 | 8.7 | 10.5 | 7.1 | 43.1 |
| Portugal | 20.5 | 14.0 | 18.3 | 8.9 | 5.1 | 8.3 | 3.7 | 21.3 |  | 11.1 | 12.0 | 17.9 | 12.8 | 8.3 | 10.4 | 2.8 | 24.6 |
| Qatar | 22.9 | 17.7 | 16.9 | 13.9 | 8.0 | 6.9 | 2.4 | 11.1 |  | 23.6 | 17.9 | 13.3 | 14.0 | 7.8 | 7.5 | 2.8 | 13.0 |
| Russian Federation | 6.7 | 11.1 | 12.7 | 15.6 | 9.6 | 8.7 | 9.0 | 26.8 |  | 6.0 | 5.1 | 11.2 | 17.3 | 11.8 | 8.7 | 7.6 | 32.4 |
| Singapore | 15.1 | 15.7 | 15.8 | 10.3 | 4.9 | 11.3 | 2.8 | 23.9 |  | 15.0 | 12.4 | 14.5 | 10.5 | 4.9 | 10.9 | 3.7 | 28.3 |
| Slovakia | 12.4 | 15.3 | 14.7 | 10.1 | 7.5 | 10.8 | 2.8 | 26.3 |  | 7.2 | 8.5 | 12.7 | 11.3 | 8.7 | 11.6 | 5.0 | 35.0 |
| Slovenia | 8.9 | 15.2 | 17.0 | 14.1 | 8.8 | 11.8 | 4.6 | 19.4 |  | 5.0 | 11.3 | 12.7 | 12.8 | 8.6 | 13.5 | 6.6 | 29.5 |
| Spain | 19.4 | 12.7 | 15.3 | 13.6 | 8.1 | 9.9 | 3.1 | 18.0 |  | 13.6 | 10.2 | 16.5 | 14.9 | 10.4 | 10.5 | 3.8 | 19.9 |
| Sweden | 11.1 | 12.3 | 13.1 | 9.9 | 8.4 | 14.3 | 6.6 | 24.2 |  | 6.6 | 7.3 | 9.4 | 11.0 | 9.0 | 14.7 | 7.9 | 34.1 |
| Switzerland | 9.0 | 15.1 | 11.7 | 11.6 | 7.3 | 11.9 | 6.0 | 27.4 |  | 6.6 | 9.4 | 11.2 | 9.2 | 7.2 | 12.4 | 7.6 | 36.4 |
| Thailand | 4.8 | 18.7 | 17.2 | 16.7 | 6.3 | 13.6 | 1.5 | 21.2 |  | 4.9 | 13.0 | 13.2 | 16.3 | 5.3 | 16.1 | 2.3 | 28.8 |
| United Arab Emirates | 22.1 | 20.5 | 16.0 | 14.8 | 7.3 | 4.4 | 4.4 | 10.5 |  | 19.4 | 20.6 | 16.5 | 12.8 | 7.6 | 4.8 | 4.4 | 14.0 |
| Tunisia | 23.3 | 22.9 | 15.6 | 10.7 | 5.7 | 7.9 | 1.1 | 12.9 |  | 13.0 | 17.1 | 14.6 | 11.2 | 7.7 | 10.9 | 2.0 | 23.4 |
| Turkey | 27.0 | 18.8 | 15.3 | 10.9 | 6.8 | 7.8 | 1.6 | 11.9 |  | 28.2 | 15.6 | 14.1 | 10.5 | 7.2 | 7.1 | 2.8 | 14.4 |
| United Kingdom | 14.3 | 17.0 | 12.5 | 11.7 | 7.3 | 11.5 | 3.6 | 22.2 |  | 8.6 | 11.3 | 13.3 | 12.9 | 8.5 | 13.5 | 5.8 | 26.1 |
| United States | 13.7 | 9.1 | 10.7 | 12.9 | 8.5 | 15.8 | 4.8 | 24.6 |  | 6.2 | 6.1 | 8.1 | 11.3 | 8.1 | 14.9 | 9.4 | 36.1 |
| Uruguay | 21.0 | 12.8 | 16.7 | 10.7 | 7.4 | 10.6 | 4.8 | 15.9 |  | 12.6 | 10.1 | 13.9 | 10.7 | 9.1 | 14.0 | 6.5 | 23.1 |

NB: percent of pupils in each category

| **(d) Out of school vigorous activity (20mins+), days/week** | | | | | | | | | | | | | | | | | |
| --- | --- | --- | --- | --- | --- | --- | --- | --- | --- | --- | --- | --- | --- | --- | --- | --- | --- |
|  | Wealth Q1 |  |  |  |  |  |  |  |  | Wealth Q5 | | |  |  |  |  |  |
| Country | 0 | 1 | 2 | 3 | 4 | 5 | 6 | 7 |  | 0 | 1 | 2 | 3 | 4 | 5 | 6 | 7 |
| Australia | 22.8 | 16.2 | 16.8 | 15.9 | 8.5 | 8.4 | 3.9 | 7.6 |  | 10.9 | 12.4 | 14.2 | 15.5 | 13.7 | 11.4 | 6.1 | 15.8 |
| Austria | 24.0 | 19.7 | 18.1 | 14.4 | 8.3 | 6.9 | 3.0 | 5.6 |  | 16.7 | 16.5 | 17.9 | 15.3 | 11.1 | 9.6 | 4.8 | 8.3 |
| Belgium | 26.0 | 21.8 | 17.0 | 13.8 | 8.8 | 5.7 | 2.4 | 4.6 |  | 12.9 | 16.4 | 17.8 | 17.9 | 10.7 | 10.3 | 4.4 | 9.6 |
| Brazil | 36.3 | 20.4 | 13.7 | 8.5 | 4.7 | 5.8 | 2.5 | 8.0 |  | 28.0 | 14.8 | 13.6 | 12.0 | 6.4 | 10.1 | 3.5 | 11.6 |
| Bulgaria | 24.3 | 19.2 | 17.3 | 13.1 | 6.3 | 6.8 | 3.8 | 9.1 |  | 12.9 | 13.4 | 16.0 | 15.7 | 10.0 | 10.9 | 5.5 | 15.6 |
| Canada | 20.2 | 14.2 | 15.2 | 14.4 | 9.7 | 13.0 | 4.1 | 9.1 |  | 9.5 | 8.0 | 13.7 | 14.1 | 12.4 | 15.1 | 7.6 | 19.8 |
| Chile | 21.3 | 23.1 | 16.6 | 14.4 | 7.4 | 7.1 | 2.5 | 7.6 |  | 19.8 | 17.9 | 16.9 | 14.0 | 9.8 | 7.5 | 3.8 | 10.3 |
| Chinese Taipei | 21.2 | 15.6 | 24.6 | 10.3 | 5.5 | 9.0 | 2.7 | 11.2 |  | 15.7 | 14.3 | 24.1 | 13.6 | 6.2 | 9.4 | 3.2 | 13.7 |
| Colombia | 21.6 | 29.3 | 15.1 | 9.4 | 7.2 | 7.5 | 2.0 | 7.9 |  | 18.5 | 20.3 | 16.1 | 12.1 | 7.7 | 10.2 | 4.6 | 10.5 |
| Costa Rica | 29.7 | 23.3 | 14.9 | 11.8 | 6.0 | 5.1 | 2.2 | 6.9 |  | 21.9 | 18.4 | 15.0 | 17.4 | 8.5 | 6.8 | 3.7 | 8.2 |
| Croatia | 24.0 | 18.7 | 17.9 | 12.5 | 8.2 | 6.1 | 3.2 | 9.4 |  | 16.5 | 13.6 | 13.9 | 15.2 | 9.5 | 10.2 | 7.7 | 13.4 |
| Czech Republic | 15.1 | 18.8 | 15.8 | 14.9 | 12.5 | 7.7 | 5.0 | 10.3 |  | 8.3 | 12.2 | 15.0 | 18.4 | 13.5 | 10.3 | 6.2 | 16.0 |
| Denmark | 13.9 | 15.3 | 16.8 | 14.7 | 11.5 | 10.2 | 5.9 | 11.7 |  | 8.7 | 8.5 | 13.2 | 18.2 | 13.5 | 14.0 | 6.8 | 17.0 |
| Dominican Republic | 16.9 | 20.8 | 15.7 | 15.0 | 8.5 | 9.7 | 3.4 | 10.0 |  | 15.0 | 15.3 | 17.4 | 12.3 | 10.1 | 10.3 | 4.9 | 14.8 |
| Estonia | 17.5 | 15.5 | 16.5 | 17.9 | 10.5 | 8.9 | 3.5 | 9.8 |  | 8.9 | 11.3 | 18.3 | 16.0 | 12.9 | 13.7 | 6.3 | 12.5 |
| Finland | 15.4 | 17.3 | 20.2 | 17.2 | 9.5 | 9.7 | 4.8 | 5.9 |  | 10.4 | 13.0 | 16.6 | 16.8 | 10.9 | 12.7 | 10.1 | 9.6 |
| France | 29.2 | 21.6 | 17.3 | 12.3 | 7.0 | 3.8 | 2.3 | 6.6 |  | 21.9 | 18.7 | 19.6 | 14.3 | 8.7 | 5.9 | 3.1 | 7.7 |
| Germany | 16.6 | 16.1 | 20.1 | 18.3 | 12.0 | 6.8 | 3.1 | 7.1 |  | 8.6 | 12.4 | 18.3 | 20.5 | 14.1 | 10.9 | 3.7 | 11.6 |
| Greece | 21.7 | 15.8 | 16.7 | 15.0 | 8.4 | 7.8 | 5.0 | 9.7 |  | 15.3 | 11.3 | 15.7 | 16.2 | 11.7 | 10.1 | 5.5 | 14.1 |
| Hong Kong | 25.7 | 25.5 | 16.3 | 10.7 | 4.8 | 6.8 | 2.8 | 7.4 |  | 18.0 | 22.6 | 17.4 | 14.3 | 6.6 | 7.4 | 2.9 | 10.9 |
| Hungary | 20.7 | 13.1 | 15.8 | 14.9 | 10.8 | 10.1 | 2.6 | 11.9 |  | 8.5 | 7.8 | 14.7 | 18.0 | 11.9 | 13.8 | 7.3 | 18.1 |
| Iceland | 10.8 | 10.1 | 13.5 | 12.1 | 9.6 | 15.7 | 9.5 | 18.6 |  | 7.7 | 7.1 | 8.6 | 10.4 | 12.1 | 15.3 | 12.6 | 26.2 |
| Ireland | 19.8 | 12.6 | 17.2 | 15.4 | 12.3 | 9.1 | 5.4 | 8.2 |  | 10.9 | 13.1 | 14.5 | 17.1 | 11.7 | 10.8 | 7.0 | 14.9 |
| Israel | 25.6 | 20.5 | 15.5 | 14.0 | 8.0 | 6.4 | 3.8 | 6.2 |  | 17.2 | 13.4 | 16.3 | 14.7 | 12.1 | 8.7 | 4.6 | 13.0 |
| Japan | 36.2 | 13.3 | 8.5 | 9.0 | 4.1 | 6.0 | 9.7 | 13.2 |  | 31.8 | 9.6 | 9.6 | 8.4 | 3.4 | 7.3 | 10.5 | 19.6 |
| Korea | 30.3 | 16.0 | 18.1 | 12.0 | 5.4 | 7.9 | 3.3 | 7.0 |  | 24.8 | 14.2 | 21.8 | 13.1 | 6.2 | 7.0 | 2.0 | 10.9 |
| Latvia | 15.7 | 15.5 | 16.5 | 17.8 | 10.7 | 10.0 | 4.7 | 9.0 |  | 8.6 | 10.2 | 17.1 | 16.3 | 13.5 | 14.6 | 5.2 | 14.5 |
| Lithuania | 18.1 | 15.9 | 19.7 | 16.2 | 9.7 | 8.8 | 3.4 | 8.1 |  | 9.7 | 10.6 | 15.3 | 16.6 | 12.4 | 13.3 | 6.9 | 15.3 |
| Luxembourg | 23.8 | 17.8 | 18.4 | 12.6 | 9.4 | 4.9 | 4.0 | 9.3 |  | 10.7 | 12.7 | 16.3 | 15.4 | 13.5 | 10.9 | 6.6 | 13.9 |
| Mexico | 14.2 | 22.7 | 21.2 | 12.4 | 7.4 | 10.6 | 2.6 | 8.9 |  | 13.2 | 13.1 | 16.5 | 12.8 | 10.8 | 15.2 | 5.5 | 13.0 |
| Montenegro | 17.5 | 16.8 | 15.2 | 15.7 | 8.4 | 8.0 | 3.8 | 14.6 |  | 11.7 | 11.0 | 13.8 | 14.6 | 9.2 | 11.3 | 7.6 | 20.8 |
| Netherlands | 25.0 | 19.0 | 15.7 | 20.2 | 9.2 | 4.7 | 2.8 | 3.3 |  | 11.2 | 13.0 | 15.0 | 24.3 | 15.6 | 9.5 | 5.3 | 6.2 |
| New Zealand | 25.9 | 13.7 | 14.9 | 11.1 | 12.3 | 9.6 | 4.8 | 7.8 |  | 12.6 | 8.7 | 17.7 | 13.8 | 12.6 | 12.0 | 7.8 | 14.9 |
| Norway | 14.3 | 14.5 | 18.0 | 16.6 | 12.4 | 10.5 | 4.4 | 9.2 |  | 7.5 | 9.3 | 11.7 | 15.0 | 15.5 | 15.0 | 9.0 | 17.0 |
| Peru | 12.0 | 29.3 | 20.9 | 11.2 | 7.7 | 7.7 | 3.5 | 7.7 |  | 17.2 | 18.4 | 17.8 | 15.2 | 8.7 | 8.4 | 4.1 | 10.2 |
| Poland | 13.4 | 13.6 | 14.9 | 16.4 | 9.2 | 10.2 | 5.3 | 17.1 |  | 10.7 | 8.7 | 11.1 | 15.1 | 11.3 | 11.4 | 8.5 | 23.2 |
| Portugal | 26.5 | 17.5 | 20.5 | 12.0 | 7.7 | 5.4 | 2.4 | 8.0 |  | 16.2 | 12.6 | 21.0 | 14.8 | 12.6 | 8.9 | 3.7 | 10.2 |
| Qatar | 26.6 | 19.4 | 17.2 | 9.9 | 7.6 | 7.6 | 2.8 | 8.8 |  | 25.3 | 17.7 | 14.7 | 12.8 | 8.7 | 7.2 | 3.9 | 9.7 |
| Russian Federation | 14.6 | 13.9 | 17.0 | 20.2 | 9.4 | 9.3 | 4.9 | 10.7 |  | 10.9 | 8.4 | 11.9 | 22.6 | 12.9 | 9.3 | 8.5 | 15.6 |
| Singapore | 19.1 | 22.4 | 22.3 | 13.6 | 8.1 | 6.3 | 1.7 | 6.5 |  | 16.7 | 21.4 | 21.2 | 15.4 | 9.6 | 7.2 | 2.8 | 5.8 |
| Slovakia | 18.1 | 17.1 | 19.4 | 15.4 | 7.7 | 7.0 | 2.8 | 12.5 |  | 9.7 | 12.9 | 15.2 | 15.7 | 10.4 | 10.4 | 6.8 | 18.9 |
| Slovenia | 17.4 | 17.0 | 16.3 | 14.3 | 9.2 | 9.8 | 4.9 | 11.1 |  | 7.9 | 11.3 | 14.0 | 13.9 | 12.4 | 11.6 | 8.9 | 20.1 |
| Spain | 24.8 | 15.0 | 19.6 | 12.2 | 11.3 | 6.1 | 3.9 | 7.0 |  | 19.6 | 10.9 | 17.8 | 17.0 | 12.7 | 10.0 | 4.7 | 7.3 |
| Sweden | 18.2 | 16.9 | 16.1 | 13.2 | 9.6 | 11.7 | 5.9 | 8.2 |  | 8.1 | 8.9 | 13.9 | 15.0 | 14.8 | 13.9 | 8.5 | 17.0 |
| Switzerland | 16.2 | 18.3 | 19.5 | 17.9 | 9.3 | 8.6 | 4.0 | 6.3 |  | 9.2 | 14.4 | 16.0 | 19.1 | 14.0 | 11.1 | 4.8 | 11.4 |
| Thailand | 10.3 | 25.0 | 23.1 | 15.5 | 6.8 | 8.2 | 1.6 | 9.5 |  | 13.8 | 21.1 | 19.7 | 16.0 | 6.7 | 9.4 | 2.4 | 11.1 |
| United Arab Emirates | 25.7 | 19.6 | 17.9 | 13.6 | 7.7 | 4.9 | 2.6 | 8.0 |  | 24.0 | 20.3 | 16.5 | 12.6 | 7.9 | 5.5 | 4.1 | 9.1 |
| Tunisia | 25.9 | 25.3 | 18.4 | 9.8 | 6.3 | 5.3 | 1.4 | 7.6 |  | 17.6 | 19.0 | 18.6 | 15.0 | 7.6 | 7.3 | 2.9 | 11.9 |
| Turkey | 26.9 | 20.8 | 15.6 | 11.4 | 7.0 | 6.2 | 2.9 | 9.3 |  | 25.5 | 19.1 | 16.1 | 10.9 | 6.1 | 8.5 | 3.5 | 10.4 |
| United Kingdom | 28.4 | 22.1 | 17.0 | 12.9 | 5.7 | 6.9 | 1.6 | 5.5 |  | 14.5 | 16.3 | 19.9 | 15.0 | 11.8 | 8.4 | 4.7 | 9.4 |
| United States | 20.1 | 12.1 | 11.0 | 13.5 | 9.2 | 14.7 | 5.5 | 13.9 |  | 12.0 | 7.6 | 10.5 | 11.0 | 9.5 | 18.4 | 9.4 | 21.7 |
| Uruguay | 32.2 | 12.4 | 17.4 | 12.4 | 6.4 | 6.0 | 4.8 | 8.4 |  | 16.6 | 10.1 | 16.7 | 17.7 | 9.7 | 11.4 | 7.3 | 10.5 |

NB: percent of pupils in each category
